# Expression profiling of the learning striatum

**DOI:** 10.1101/2023.01.03.522560

**Authors:** E Lousada, Z Kliesmete, A Janjic, D Richter, LE Wange, E Briem, I Hellmann, E Burguière, W Enard, C Schreiweis

**Affiliations:** Sorbonne Université, Institut du Cerveau - Paris Brain Institute - ICM, Inserm, CNRS, AP-HP, Hôpital de la Pitié Salpêtrière, Paris, France; Anthropology and Human Genomics, Faculty of Biology, Ludwig-Maximilians-Universität München, Germany; CNRS, INSERM, Centre de Recherche en Neurosciences de Lyon CRNL U1028, UMR5292, Team Neurophysiology of Repetitive Behaviours, Université Claude Bernard Lyon 1, Centre Hospitalier Le Vinatier, Bron, France; Université Claude Bernard Lyon 1 (UCBL), 69622 Villeurbanne Cedex, France

## Abstract

How learning reshapes the brain across temporal and spatial scales, from molecular to cellular levels, remains a fundamental question in neuroscience. In the striatum, ventromedial, dorsomedial and dorsolateral subregions are differentially recruited during early to late learning on a neurophysiological level. Here, we profile the transcriptome of these subregions across 396 biopsies from 66 mice to map molecular dynamics at three learning stages of a visual discrimination task. We find a strong transcriptional change during the early phase that diminishes towards later learning and is indistinguishable among subregions. The 818 genes altered during learning are enriched for neuronal processes, but also include circadian, vascular, and oligodendrocyte-related pathways. We provide a web application for this large learning-related expression dataset, enabling interactive exploration of genes, gene sets, and transcription factor activity, and suggest that this framework will be broadly useful for diverse neuroscience questions.

## Introduction

Understanding the mechanisms underlying learning is a fundamental goal in neuroscience. Whether foraging for food, playing the piano, or driving a car, learning and mastering such capacities rely on the acquisition and repeated execution of action sequences. Motivation drives the initial stage of learning, encouraging high amounts of exploration and trial-and-error^1,2^. As learning progresses, associations between sensory cues, specific actions, and their consequences increase the efficiency of the behaviour^1,3^. With sufficient repetition, behaviour tends to shift from explorative to exploitative strategies, and get partially automatized^4–6^. Cortico-basal ganglia circuits are crucial for this type of learning. Classically, the cortico-basal ganglia architecture is organized into three different loops that are differentially recruited throughout different learning stages. During early learning, the limbic and associative loops are thought to guide exploration and goal-directed action selection^7,8^. With extended practice, the sensorimotor loops become increasingly engaged, supporting efficient and automatized execution of well-learned action sequences^9,8,10^.

The striatum, the major input nucleus of the basal ganglia, mirrors this compartmentalized organization. In rodents, the limbic, associative and sensorimotor loops course through ventromedial (VMS), dorsomedial (DMS) and dorsolateral (DLS) territories, respectively^11,12^. Although this tripartite scheme is an oversimplification, and more fine-grained distinctions have been proposed based on extensive anatomical and physiological work^13,14^, it remains a useful framework for describing the topographic organisation of the striatum and its cortical connections. The striatum is primarily composed of inhibitory, GABAergic medium spiny neurons (MSNs), which constitute over 95% of the neuronal population and project onto the output basal ganglia nuclei. These MSNs are modulated by dense excitatory, glutamatergic inputs from the cortex and thalamus, by a variety of interneurons (both GABAergic and cholinergic), as well as by dopaminergic afferents from the midbrain, all of which seem to have an essential role in reward-related learning, motivation, and plasticity^15–18^.

Learning leaves enduring neuroplastic traces as a neural engram in involved circuitry, ranging from synaptic strengthening or remodeling^19,20^ to functional connectivity changes between different brain regions^21,22^. More recently, the importance of non-neuronal cells in learning and circuit plasticity has also become increasingly evident^23^. Ultimately, learning relies on coordinated molecular changes that reconfigure neuronal and non-neuronal function. Large-scale, single-cell transcriptomic approaches have mapped the molecular signatures of the different cell types in the striatum^24,25^ including a recent in-depth analysis of over a million single-cell profiles from murine GABAergic neurons in the adult and developing telencephalon^26^. Such atlases provide an invaluable catalogue of cell types and states with high spatial and temporal resolution. However, how learning affects the gene expression programs of these cells remains largely unexplored, as this requires many biological replicates and hence efficient behavioural and expression readouts.

Here, we combine an automated behavioural assay with efficient bulk RNA-seq to address this gap with reasonable statistical power. We collected a total of 396 expression profiles from punch biopsies of the ventral, dorsomedial and dorsolateral striatum of a total of 66 mice during early, intermediate and late stages of learning, and their non-learning matched controls. Our experimental design and large number of biological replicates enable the first quantitative comparison of gene expression changes across three striatal regions and learning stages, revealing a strong response during early learning that is largely region-unspecific. The 818 identified genes that change expression during learning indicate a variety of neuronal (e.g. synaptic plasticity) and non-neuronal (e.g. angiogenesis) cellular responses. To our knowledge, this represents the largest dataset linking learning behavior to gene expression signatures, and we provide an interactive web application for exploring this expression atlas of the learning striatum (https://shiny.bio.lmu.de/Dopaloops/).

## Materials and Methods

### Animals

All experimental procedures followed national and European guidelines and were approved by the institutional review boards (French Ministry of Higher Education, Research and Innovation; APAFiS Protocols no. 1418-2015120217347265 and 2021042017105235). Animals were group-housed in the animal facilities of the Paris Brain Institute in Tecniplast ventilated polycarbonate cages under positive pressure with hard-wood bedding in groups of up to six animals per cage with *ad libitum* food and water access. The temperature was maintained at 21–23 °C and the relative humidity at 55 ± 10% with a 12-h light/dark cycle (lights on/off at 8am and 8pm, respectively). Following ethical guidelines of animal experimentation on sex balance and the principle of the three Rs, we included both male and female adult mice (aged 5.50 ± 1.88 months). The behavioural task was validated in a total of 25 wildtype animals (N = 19 males and N = 6 females) on C57BL6/J background. For the RNA-sequencing experiment, a total number of N = 66 wildtype animals on C57BL6/J background were used (N = 50 males and N = 16 females).

### Experimental setup and task

Operant conditioning was conducted in custom-modified experimental chambers (ENV-007CTX, Med Associates, Vermont, USA) as previously described^27^, in which each of the tested animals lives and performs the conditioning task 24/7 in an automated, self-initiated, self-paced manner. Seven such experimental operant chambers were operated in parallel in the same experimental room (Figure 1A). Briefly, the rear wall of each chamber housed the feeder compartment (equipped with an infrared light beam to detect feeder entry crossing) and drinking water bottle holder, the front wall held two tactile screens, and a custom-made pair of black Plexiglas gates equipped with a pair of infrared (IR) beams. These beams allowed for tracking the locomotion of the mouse into or out of the area with the tactile screens (Figure 1B-C). The chambers of the yoked controls were custom-built, plexiglass chambers as described in Lamothe et al., 2023^28^ and were located in the same experimental room as the operant conditioning chambers. The yoked control chambers contained *ad libitum* water access as well as the same woodchip bedding and cotton pad nesting material as the operant conditioning chambers and regular housing cages in the facilities. Precision reward pellets served as the sole nutrition during the entire experiment. The reward pellets (20 mg LabTabTM AIN-76A rodent precision pellets, TestDiet, Richmond, USA) were earned upon successful trials by the mice in the conditioning task (“learners”); yoked control mice, matched for sex, age and whenever possible also for litter, received the exact same amount of food rewards as their matched pair undergoing actual conditioning.

**Figure 1.**
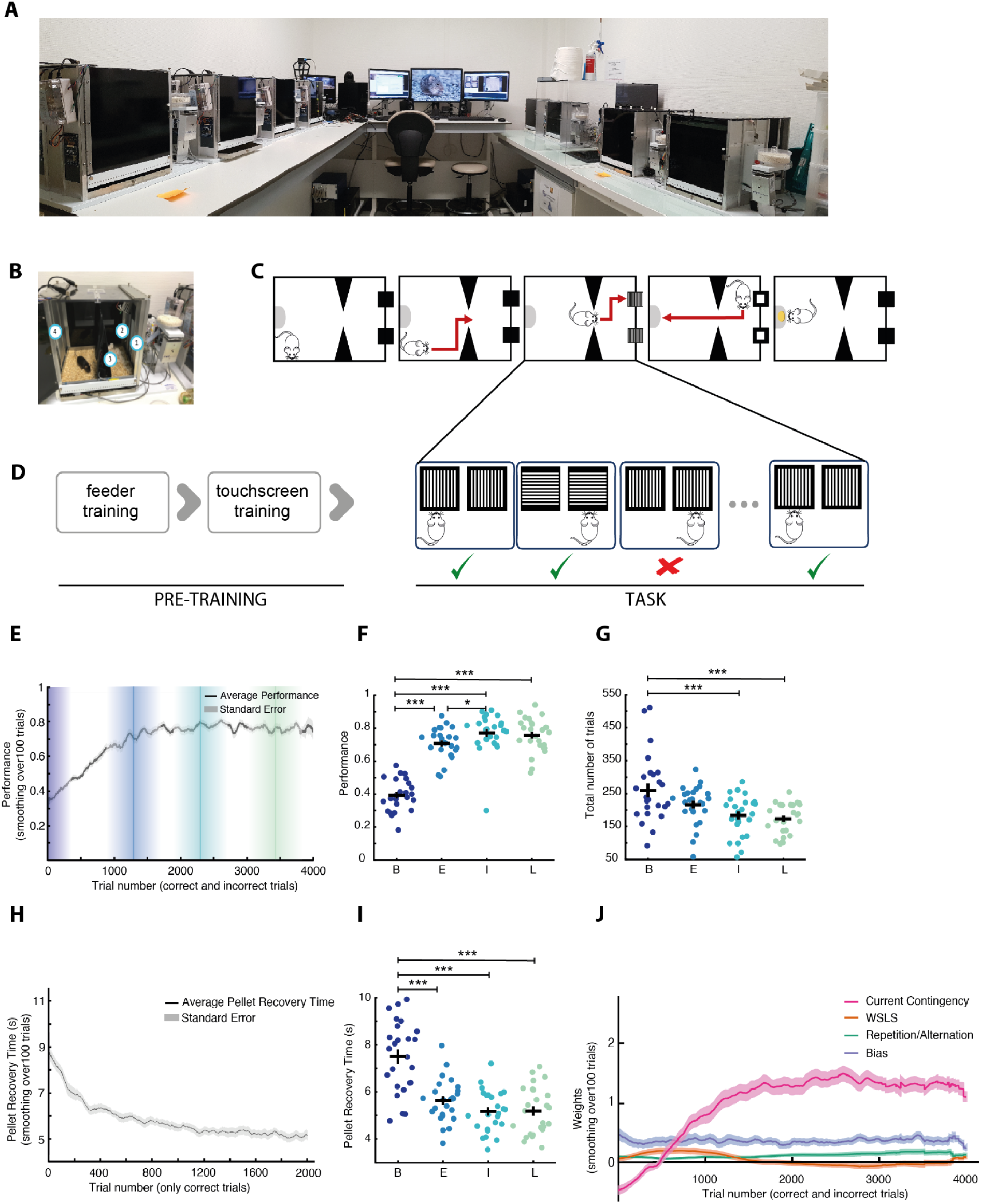
An automated visual discrimination task for associative action-outcome learning. A – Photo of the behavioural apparatus with seven operant conditioning chambers running at the same time. B – Close-up photo of one of the operant conditioning chambers. 1 - feeder, 2 - water dispenser, 3 - tunnel, 4 - screens. C – Schematic of the custom-built operant conditioning chamber (top view). Mice live in the chamber continuously and self-initiate trials by crossing an infrared beam-equipped gate. Trials present one of two visual stimuli (vertical or horizontal bars) on dual touchscreens. A correct response triggers reward delivery at the rear food dispenser. D – Overview of task structure. Following a pre-training phase (feeder and touchscreen familiarization), mice enter a binary visual discrimination task where they must associate bar orientation (vertical or horizontal) with a specific touchscreen response to obtain a food pellet. E – Performance follows a sigmoidal trajectory over the course of learning, reaching a plateau at ∼1329 trials. Shown is the mean ± SEM across 25 wild-type mice. Shaded regions represent the four defined learning stages: baseline (dark blue), early (blue), intermediate (turquoise), and late (green). F – Individual-level data across learning stages for performance. Significant improvements between baseline and early, intermediate, and late stages; and between early and intermediate. G – Individual-level data across learning stages for the number of initiated trials (completed and non-completed). Significant improvements between baseline and intermediate, and late stages. H – Pellet recovery time (PRT), defined as the latency between correct screen touch and reward retrieval, decreases markedly with the learning stage, reflecting increased behavioural efficiency. Values represent mean ± SEM. I – Individual-level data across learning stages for PRT. Significant improvements between baseline and early, intermediate, and late stages. J – Latent decision strategies estimated using PsyTrack. Strategy weights over time reveal a progressive increase in reliance on the correct stimulus–response rule (*current contingency*, pink), which closely follows the performance curve. Other strategies: *WSLS* (orange), *repetition/alternation* (green), and *lateralization bias* (violet) remain stable or decline over time. Data in F, G, I shown as means ± SEM; color code consistent across panels. B: baseline, E: early, I: intermediate, L: late. Significance levels: ***p < 0.001; **p < 0.01; *p < 0.05.

Once placed in the operant conditioning chambers, the mice started with a pre-training phase of four different stages. In a first stage, a total of five precision pellets dropped automatically once every minute if the previous pellet had successfully been recovered. In a second stage, the mouse was required to nose-poke into the feeder compartment to trigger individual pellet delivery once per minute until a total of 10 pellets had been received. In a third stage, crossing of the IR photobeams prompted the illumination of both tactile screens; the mouse was required to touch one of them to trigger pellet delivery in the feeder. There was no time limit for recovering the pellets at this stage. However, the condition for the mice to pass into the following pre-training stage was the recovery of the pellets within less than 15 seconds for at least 66% of the times. Then, in the consecutive pre-training stage, as before, both screens were illuminated with white light upon gate crossing. However, mice now had to nose-poke into the feeder within less than 15 seconds after screen touch in order to trigger pellet delivery. Having successfully recovered at least 66% of the pellets in this manner, mice passed into the actual task.

The behavioural task was a deterministic, binary visual discrimination task, where mice self-initiated each trial by triggering visual stimulus onset on the tactile screens by again crossing the photobeam-equipped gate. The presented stimulus pair consists of ten white, either vertical or horizontal, bars on a black screen, each of which was presented 50% of the time in a random manner (Figure 1D). To earn a precision pellet, mice needed to associate each of the two stimuli with one screen touch, e.g. stimulus of ten horizontal/ vertical bars associated with the action of left/ right screen touch, accordingly. The contingencies were balanced across individuals and operant chambers. Mice were allowed a maximum of 60 seconds after stimulus onset to respond with a screen touch. Upon correct response, the screens started blinking to give positive feedback for the mouse, which had 15 seconds to nose poke in the food dispenser in the rear wall for obtaining a food reward (Figure 1C).

Past 15 seconds after the correct response, the trial became unrewarded. In case of incorrect screen touch, mice received an aversive white light for five seconds. Additionally, in order to decrease the tendency of mice to lateralize, each trial following an incorrect response required the mouse to switch to the other screen. Past 60 seconds in the chamber compartment with the screens, the screens would switch off again; in these time out trials, mice then had to cross out and back into the feeder compartment to trigger the next trial. Trials where mice crossed in and out of the screen compartment within 60 seconds and without screen touch, were quantified aborted trials (Extended Data Figure S1). Animals for validation of the task and subsequent determination of the time points of baseline, early, intermediate and late stages of learning remained in the task for around 5,500 trials (trial number = 5,517 +/- 1,177; mean +/- SD, respectively). Baseline time point was defined as the first complete night after pre-training, when animals reached the task stage. The time point of the early learning stage was estimated at the individual-subject level based on the onset of sustained high performance during learning. For each subject, performance trajectories were smoothed using a sliding moving average (window = 150 trials) to reduce trial-to-trial variability while preserving learning dynamics. A performance plateau was defined as the mean of the top 20% of smoothed performance values. The early learning time point was operationally defined as the first trial initiating a contiguous block of at least 300 trials for which smoothed performance remained within Δ = 0.10 of the estimated plateau level, allowing for normal behavioural variability while ensuring sustained performance rather than transient fluctuations. In cases where no block met these criteria, the time point was conservatively set to the trial corresponding to maximal smoothed performance. Detection parameters were chosen to balance sensitivity and specificity and were validated by visual inspection of all individual learning curves. Using this procedure, the early learning time point occurred at a mean trial number of 2,728.8 +/- 786.5 (mean +/- SD, respectively) across subjects. If we remove the non-completed trials, and thus including only the trials where the subject performed a correct or an incorrect response, the equivalent in trial number is 1,328.8 ± 439.2 (mean +/- SD, respectively) (Extended Data Figure S2). Intermediate and late learning time points were determined as 1,000 and 2,000 trials post early learning stage time point, respectively.

In the RNA-seq cohort of mice, we could not define the learning stage from a later performance curve analysis. For this reason, and to account for inter-individual variability and ensure we captured transitional processes, the learning stages were defined slightly earlier than the average inflection points observed in the validation cohort, corresponding to 1,000, 2,000, and 3,000 completed trials, respectively. To control for experimental factors (e.g. social isolation and food consumption), each learning mouse was paired with a matched animal of the same genotype, sex, and age as a yoked control. Learners and yoked controls were put into their according behavioural setup at the same time, and sacrificed at the same time, with a randomized order of the dissection/tissue harvesting being randomized for learning condition. The yoked controls received the exact same amount of precision food pellets as their learning partners during the past 24 hours in the mornings.

### Sample collection

Having reached respective criteria for early/intermediate/late learning stages, the brains of both learners and their paired yoked partners were rapidly dissected and collected during the morning of the following day. Before each dissection, all tools and surfaces were cleaned with RNAse decontamination solution (Fisher Scientific, reference #10180601). Animals were euthanized by cervical dislocation, rapidly dissected and their brains immediately collected using a 5-blade custom-made tool to rapidly obtain coronal slices of 1mm thickness at similar locations across mice. All dissections were performed in a petri dish, which had been cleaned with RNAse decontamination solution beforehand. Brains were sprinkled with sterile saline to remove excess blood. All samples were collected by the same experienced researcher using 1mm^2^ biopsy punchers (pfm medical, reference #49101) to collect 1mm^3^ samples from each hemisphere of the three regions of interest: ventromedial striatum (AP = 1.10, ML = ±1.00, DV = 4.00), dorsomedial striatum (AP = 0.14, ML = ±1.50, DV = 2.10), and dorsolateral striatum (AP = 0.14, ML = ±2.50, DV = 2.50). Samples were rapidly placed inside of an RNAse-free Eppendorf tube (Dominique Dutscher, reference #033872) containing 200µL of buffer RLT Plus (Qiagen, reference #1053393), and immediately frozen on dry ice. All samples were transferred and kept at -80°C until shipment, which was done on dry ice.

### Expression data generation

Samples were transferred to 96-well plates and RNA purification and RNA-sequencing were carried out with 50µl lysate. RNA-sequencing was conducted using the prime-seq method developed by Janjic et al., 2022^29^ or updated by Pförtner et al., 2025^30^. The complete protocols for prime-seq, including primer sequences, are available, respectively, at protocols.io: (https://www.protocols.io/view/prime-seq-81wgb1pw3vpk/v2), (https://www.protocols.io/view/prime-seq-2-14egn97kpl5d/v1)

Briefly, cDNA synthesis was performed using Maxima H Minus reverse transcriptase, oligo-dT primer E3V7NEXT, and template switching oligo. After pooling, remaining primers were removed with Exonuclease I. Subsequently, cDNAs were pre-amplified using KAPA HiFi HotStart polymerase. Libraries were prepared using the NEBNext Ultra II FS DNA Library Prep Kit for Illumina (New England Biolabs, E7805S) with a custom ligation adapter, and dual indexing with TruSeq i5 and Nextera i7 index primers. Alternatively, for a second batch, RNA was extracted following the prime-seq 2 protocol. RNAs were quantified (Promega, E3310) normalized to 1 ng/µl and 4 ng RNA per sample were used for reverse transcription, pooling and cDNA amplification. Library concentrations were quantified before sequencing utilising an Agilent 2100 Bioanalyzer instrument.

Multiple rounds of sequencing were performed for two batches of learners and their respective controls. The first library was generated using Illumina NextSeq 2000 flowcell with Read 1: 28 bp (barcode: 12 bp, UMI: 16 bp), Index 1 (i7): 8 bp, Index 2 (i5): 8 bases, Read 2 (cDNA): 100 bp. The second library was sequenced on 4 Illumina NextSeq 2000 P3 flowcells and as a spike-in on a NovaSeq 6000 with the following layout: Read 1: 28 cycles (barcode: 12 cycles, UMI: 16 cycles), Index 1 (i7): 6 cycles, Index 2 (i5): 6 cycles, Read 2 (cDNA): 48 cycles (or 93 cycles on the NovaSeq Flowcell).

### Data analysis

#### Behavioural analyses

All behavioural data were analysed using Matlab R2025a. For behavioural strategy analyses, we additionally applied the PsyTrack algorithm^31^. Statistical analysis was performed using R version 4.5.2. A significance threshold of p < 0.05 was used for all analyses. We used generalized linear mixed models (GLMM) to assess the effects of learning stage, sex, and genotype on four behavioural measures: performance, number of trials, response time (RT) and pellet recovery time (PRT). Mouse identity was included as a random intercept to account for repeated measurements within individuals. For performance, RT, and PRT, models were fitted assuming Gaussian error distributions, whereas the number of trials was modelled using a count-based, negative binomial distribution. The general model structure was:

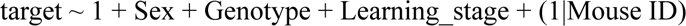

Age at start, cage, and batch were initially considered but removed after preliminary analysis indicated they did not improve model fit and contributed to convergence issues. Post-hoc comparisons between learning stages were performed using estimated marginal means (emmeans), with p-values corrected for multiple comparisons using the Holm method.

In the RNA-sequencing cohort, we applied Generalized Linear Models (GLM) instead of GLMM, since these groups consisted of independent animals rather than repeated measures within individuals. The GLM models were implemented as follows:

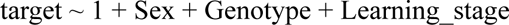

We selected a Gaussian GLM for performance, RT, and PRT due to their continuous distributions. Trial counts, being discrete and non-negative, were modeled using a negative binomial GLM to better capture the distributional characteristics of these data. We confirmed model assumptions and conducted post-hoc comparisons using estimated marginal means (emmeans) with p-values corrected for multiple comparisons using the Holm method. These analyses ensured consistency between learning cohorts and allowed for robust validation of the behavioural trends across learning stages.

#### Expression data processing and mapping

Fastq files were trimmed to remove the poly(A) tail using Cutadapt v4.1^32^. The quality was assessed using fastqc v.0.11.5^33^. zUMIs pipeline v.2.9.4d^34^ was used to process the data and generate a count matrix. Reads with a Phred quality score threshold of 20 for 3 BC bases and 4 UMI bases were filtered, mapped to the mouse genome (GRCm38) with the Gencode annotation (vM25) using STAR v.2.7.3a^35^, and then counted using RSubread v.1.32.4^36^. Genes that were expressed in at least 25% of the samples with an average of 3 counts or higher across samples from at least one striatal region were kept for the downstream analyses. This resulted in a count matrix containing 16,880 genes across 396 samples.

#### Cell-type decomposition

SCDC v.0.0.0.9000^37^ was used for cell-type deconvolution, using the scRNA-seq TM_Brain_Non_Myeloid striatal cell reference from Tabula Muris^38^. We observed high and slightly varying proportions of inhibitory neurons (80 - 96%) across samples. These proportions are likely variable due to technical reasons, as we did not observe any systematic difference in overall cell-type composition between mouse samples under learning conditions vs controls (multinomial test p-value=1.0).

#### Differential expression analysis

The input expression matrix was normalized using scaling factors calculated via TMM method from R package edgeR v.4.2.2^39^. Differential expression (DE) analysis was performed using DREAM^40^, R package variancePartition v.1.34.0^41^, which implements mixed-effect modelling. In addition to the main effects of interest, Condition and Region, we also included other nuisance factors into the following main model (model 1):

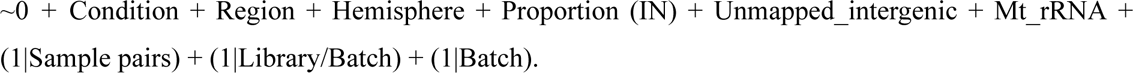

The importance of the included predictors was validated using the variance partition implemented in variancePartition as a part of the DREAM workflow. Benjamini-Hochberg adjusted p-value < 0.05 was considered as a cut-off for significant DE. To estimate the dependence of the estimated effect size and the number of detected DE genes on individual mouse pairs, we used jackknifing, e.g., systematically removed each mouse pair and refitted the model in DREAM. The distribution of estimates was summarized by the 95% central percentile interval (PI), defined by the 2.5th and 97.5th percentiles.

Residuals were obtained using the function fitVarPartModel implemented in variancePartition. Rtsne v.0.17 package with perplexity = 20 and theta = 0.1 on the top 50 PCs was used for dimension reduction during exploratory analyses.

To test whether there are genes that show significant difference in the response to learning across striatal regions, we fitted the above-mentioned model while including the interaction term Region:Condition (model 2):

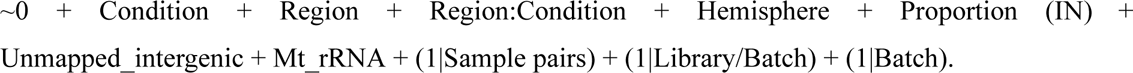

No gene passed the significance threshold of adjusted p-value 0.05. To further assess whether the learning effects are indeed similar across striatal regions, we subsetted the count matrix to each region separately and fitted the following model (model 3):

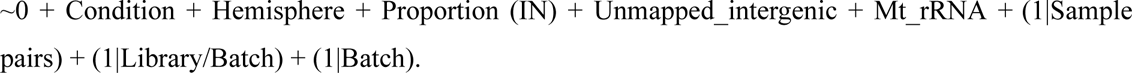

Again, jackknifing was performed where each mouse pair was systematically removed and the model refitted.

#### Gene set enrichment analysis (GSEA)

Gene Ontology analysis was performed using R package topGO v.2.56.0^42^ category Biological Process (BP), using elimination algorithm and setting minimal node size to 15. Enriched terms (Fisher’s p-value < 0.01) were further filtered for terms with <300 annotated genes and >2 significant genes to facilitate interpretability. We then sort first by p-value and then by enrichment and display the top 5 categories for a particular comparison. We also performed GSEA on MSigDb v.7.5.1 non-hierarchical annotations^43,44^. Specifically, we used collections M2 (Curated gene sets, subsetted to 3646 genes belonging to 77 sets) and M8 (Cell type signature gene sets, subsetted to 1550 genes belonging to 9 sets). Benjamini-Hochberg adjusted Fisher’s test p-value was used to identify significant enrichment, using a cutoff of 0.05.

#### Transcription factor activity scoring

Differential transcription factor (TF) activity between groups in a contrast was obtained using annotations from CollecTRI^45^ using decoupleR v.2.10.0^46^. Regulatory activities were inferred with the weighted mean method (wmean). For each contrast, the standardised gene-level z-statistic from DE analyses was used as input, together with a signed TF–target network annotation for mouse provided by CollecTRI. Wmean combines gene-level statistics across each TF’s target genes, weighted by the sign of regulation, to estimate TF activity scores. CollecTRI contains TF-target annotations for 1072 mouse TFs, 663 of which were expressed in our data. Regulators represented by fewer than 5 targets in the dataset were further excluded. Empirical significance was estimated from 100 permutations, keeping only differential TF activities between conditions (p-value<0.05).

Strong correlation observed between the TF and target expression was further used to narrow down important interactions present in the data. We used the residual count matrix, corrected for all predictors in model 1 (Section Differential expression analysis) except for Condition as the input for calculating TF-target and TF-background gene correlations. Then, one-sided Wilcox rank sum test was used to test whether the absolute correlation coefficients of annotated targets of the TF in question are higher than that of other genes with the same TF (e.g., comparison between absolute TF-target correlation and TF-background gene correlation) (p-value<0.05).

## Results

### An automated visual discrimination task for associative action-outcome learning

To investigate the transcriptional programs underlying CBG-dependent learning, we required a behavioural paradigm that allows for precise temporal staging across a large volume of trials while minimizing experimenter interference. Classical tasks such as lever-press or T-maze paradigms rely on manual training and restricted access schedules, potentially constraining the natural dynamics of learning and limiting experimental throughput. To overcome these constraints and to better approximate naturalistic behaviour, we developed a fully automated, self-motivated, visual discrimination task. Mice continuously learn a deterministic stimulus–outcome contingency in a homecage-like environment, self-initiating trials by crossing an infrared beam–equipped gate (Figure 1A-D). This triggered the display of one of two possible visual stimuli (either ten white vertical or ten horizontal bars on a black background) simultaneously on both of the two touchscreens facing the gate. Each mouse (N = 25) was assigned one of the two possible stimulus-outcome contingencies, which were balanced across mice (e.g. touching the left/right screen upon appearance of vertical/horizontal bars, respectively). Each correct response was rewarded with a small food pellet, delivered upon nose-poking into the reward feeder. We evaluated all trials during the animals’ wake cycle (8pm-8am) and distinguished between completed trials (correct or incorrect) and trials that remained non-completed for different reasons and that were classified accordingly (aborted, time-out, pellet non-retrieved, see Extended Data Figure S1 and Materials and Methods section for detailed categorization). Mice successfully acquired the task, exhibiting a typical learning curve with steadily increasing performance (correct trials / completed trials) that plateaued at 1,329 ± 439 completed trials (mean ± standard deviation; Figure 1E, Extended Data Figure S2). We defined four learning stages: baseline, early, intermediate, and late. Baseline corresponded to the first night after successfully finishing the pre-training phase, i.e. mice had successfully acquired the interaction of touching any of the two touchscreens followed by nose-poking in the feeder in order to trigger dispensing of the food pellet reward. Early learning corresponded to the night on which performance reached a plateau (Extended Data Figure S2); intermediate, to the night 1,000 trials after the performance plateau; and late, to the night 2,000 trials after the plateau (see Materials and Methods for detailed description).

In addition to performance, we quantified learning progression using the number of initiated trials (all events of crossing the infrared beam–equipped gate, i.e. all complete and non-completed trials), response time (time from stimulus onset to screen-touch) and pellet recovery time (time from screen-touch to poking into the reward feeder). All parameters were tested using a generalized linear mixed model (GLMM) with learning stage, mouse line and sex as fixed effects and Mouse ID as a random intercept (Extended Data Table 1). As expected, performance varied as a function of the learning stage. Relative to baseline, correct responses significantly increased in all time points (all Holm-corrected *p* < 0.001). Furthermore, performance significantly increased from early to intermediate time point (Holm-corrected *p* < 0.05), and stabilised after (Holm-corrected *p* > 0.05) (Figure 1F; Extended Data Table 1). The total number of initiated trials tended to decrease over task duration, with intermediate and late time points significantly dropping when compared to the baseline (Holm corrected *p* < 0.001; Figure 1G; Extended Data Table 1). Pellet recovery time dropped significantly compared to baseline across all learning stages (all Holm-corrected *p* < 0.001; Figure 1H-I; Extended Data Table 1). This indicates that mice rapidly and consistently learned to associate the stimulus with the required action, and how to get the desired reward efficiently. Response time exhibited variability across animals and learning stages (Extended Data Figure S3; Extended Data Table 1; Holm-corrected *p* > 0.05).

To further characterize behavioural strategies in the task, we used the PsyTrack model^31^ and focused on four complementary strategies that mice could apply to solve task trials: *current contingency* (i.e. following the stimulus to earn the associated outcome), *win-stay-lose-shift (WSLS;* i.e. repeat a rewarded action and switch after a non-rewarded one), *repetition/alternation* (i.e. tendency to often repeat, or often alternate the previous action) and *lateralization bias* (i.e. preferring one touchscreen). As expected from the improvement in performance described above, the weight of the *current contingency* strategy progressively increased over the course of learning, while lateralization bias and the application of a repetition/alternation strategy remained stable throughout the task. The contribution of the *WSLS* strategy transiently increased early in learning and then diminished as animals adopted the optimal, *current contingency* strategy (Figure 1J).

Altogether, these results demonstrate that the task provides a robust and powerful readout of learning, capturing both early acquisition dynamics, performance stabilization as assessed through behavioural measures of performance as well as hidden learning strategy analyses.

### Generating striatal expression profiles from learning mice and controls

To profile gene expression at different learning stages, we used the same behavioural setup as described above. Early, intermediate and late learning stages were defined using fixed thresholds of 1,000, 2,000 and 3,000 completed trials after training onset, respectively, as a behavior-based definition following tissue collection is not possible. These numbers of trials were reached on average after 7, 13 and 21 days. Crucially, each learning mouse was paired with an age- and sex-matched “yoked” control that remained in a control chamber located in the same experimental room and received an identical number of reward pellets (Figure 2A). When a learning mouse reached the required number of trials for its assigned learning stage, the learner and its yoked control were sacrificed the following morning in randomized order. Brains were sliced into 1-mm sections, and 1-mm³ punches were collected from the ventromedial (VMS), dorsomedial (DMS), and dorsolateral (DLS) striatum of each hemisphere. We ran multiple batches to obtain a total of eleven mice per group (total N = 66; Figure 2A). Behavioural measures and learning progression matched those reported for the cohort described above (Figure 2B, Extended Data Figure S4, Extended Data Table 2).

**Figure 2.**
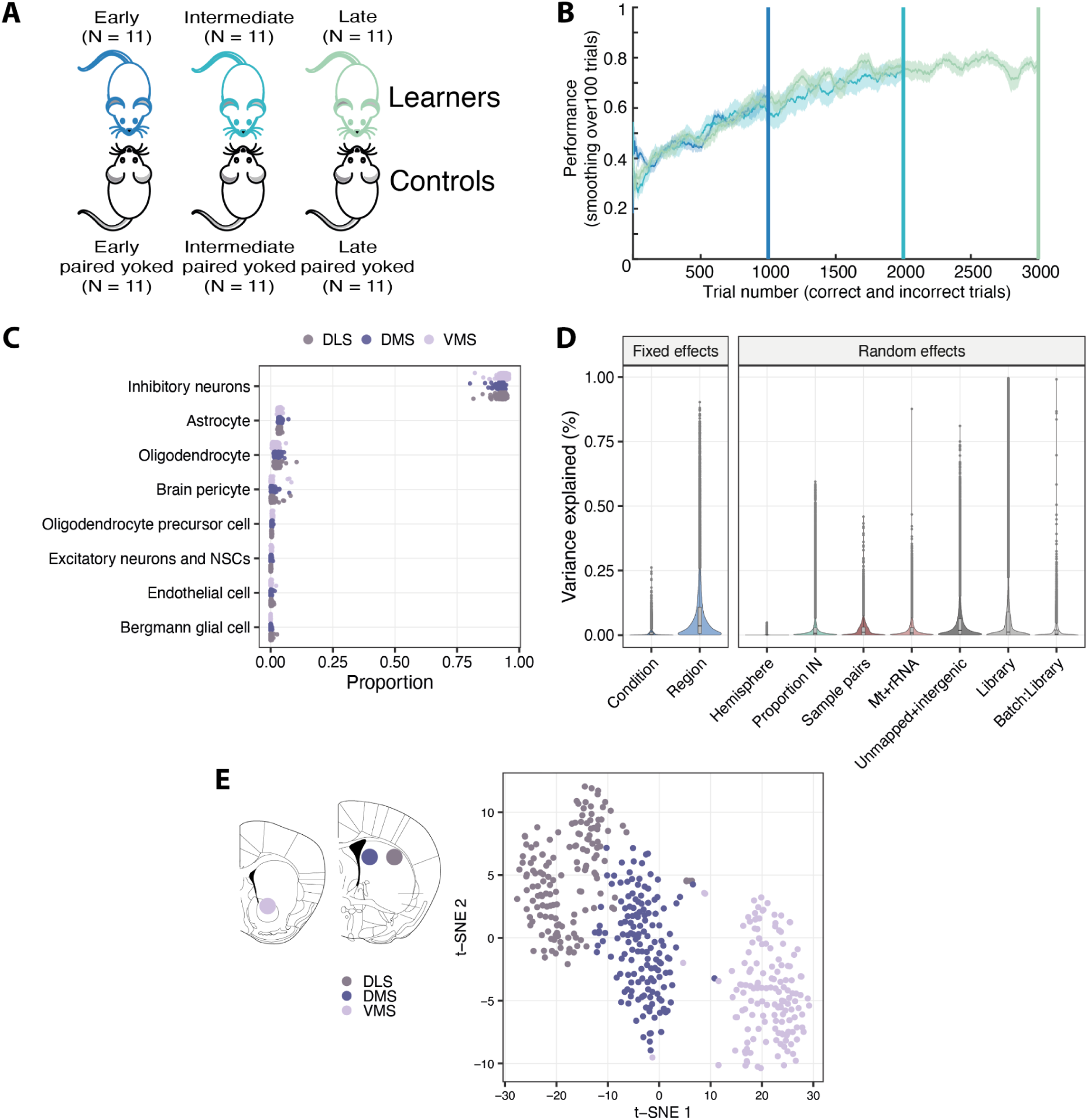
Striatal expression profiles from learning mice and controls. A – Illustration of the experimental design. Learners were paired with yoked controls to account for non-task-related variables. B – Mean performance (± SEM) of each learning stage group up to the respective stage criterion: 1000, 2000, and 3000 completed trials for early, intermediate, and late learners, respectively. Coloured lines represent the three defined learning stages: early (blue), intermediate (turquoise), and late (green). C – Cell type decomposition across samples, per region, using single-cell RNA-seq from mouse striatum (Tabula Muris) as the reference. Each dot represents a sample. The majority of the cells within the bulk samples are inhibitory neurons (80-96%). No systematic differences in the cell type composition were observed between learners and yoked mice (multinomial p-value 1.0). D – Variance partition using the predictors included in the mixed-effects modelling of gene expression using DREAM. Each dot represents a gene. The striatal region explains a large proportion of the expression variance across samples (7078 genes with >=5%). The learning condition explains at least 5% of the variance for 304 genes. E – Left: schematic overview of the three striatal regions included in the study. (VMS: AP = 1.10, ML = ±1.00, DV = 4.00; DMS: AP = 0.14, ML = ±1.50, DV = 2.10; DLS: AP = 0.14, ML = ±2.50, DV = 2.50). VMS is depicted in light purple, DMS in dark blue, DLS in grey. Right: Dimensionality reduction using t-SNE based on the 1000 most variable genes (number of PCs = 50, perplexity = 20, theta = 0.1). All random effect predictors (see methods) have been regressed out prior to clustering.

We then processed the resulting 396 punch biopsies (66 mice × 3 regions × 2 hemispheres) using prime-seq, a bulk RNA-seq method that combines bead-based RNA isolation and early barcoding of oligo(dT)-primed cDNA^29^. Its cost-efficiency (∼$3.50 per library) enabled this experimental design with many libraries, and its sensitivity allowed us to use one quarter of each 1 mm³ biopsy per library. To draft how many and which cells are expected to contribute to such an expression profile, we can make the following ballpark estimates: In the C57BL/6J striatum there are ∼73,000 neurons/mm³ and these make up 70%-80% of all cells^47^. 97 % of neurons are medium spiny neurons^47^ and the rest various interneuron types^48^. Non-neural cells are composed roughly of equal proportions of astrocytes, microglia, and oligodendrocytes^49^. Hence, each RNA-seq profile is expected to contain RNA from 18,000 medium spiny neurons, 1,000 interneurons, 2,000 astrocytes, 2,000 microglia, and 2,000 oligodendrocytes. Hence, the majority of the signal is derived from MSNs and even lowly expressed genes (e.g. 1 transcript per 100,000 transcripts in a MSN) are expected to have 7 reads per million reads in one RNA-seq profile. The sensitivity is proportionally lower for the less abundant cell types, but the statistical power to detect differently expressed genes depends also on many other factors, especially on the number of independent replicates, the magnitude of expression change and the fraction of cells in which the expression change occurs. We do not know how independent the expression measurements within regions and hemispheres of the same mouse are and we know even less about the magnitude and cellular distribution of learning-related expression changes. Nevertheless, it might be fair to say that we have reasonable power (e.g. >80%) for a reasonably well expressed gene (e.g. >3 transcripts per 100,000) that shows a reasonably big change (e.g. >2-fold upregulation) in a reasonable fraction of cells (e.g. >10%) when comparing two striatal regions of 11 mice each, given a total of 1 million unique reads (UMIs) for each sample. We generated a mean of 1.5 million unique reads (UMIs) for each of the 396 samples, to be on the safe side regarding these calculations. To our knowledge this represents the largest dataset that links a learning behaviour with gene expression signatures.

### Striatal expression profiles from learning mice and controls

We mapped 677 million reads to a total of 22,400 genes. Lowly expressed genes with limited power to detect differential expression were removed, retaining 16,880 genes that were expressed in at least 25% of the samples from at least one striatal region with an average of at least three UMIs. To estimate from which cell types these profiles were generated, we deconvoluted the expression profiles^37^ using single-cell RNA-seq data from annotated striatal cell types^38^. As expected according to striatal cell type composition (see above), the great majority of cells in our samples were inhibitory (80-96%), followed by astrocytes (2-7%) and oligodendrocytes (0.1-10%) (Figure 2C). The proportion of inhibitory cells was a good predictor for variations in other cell types and did not show any association with the striatal region or learning stage (see Materials and Methods). Hence, we interpret this variation as arising from variability in sampling (e.g., punching a bit of corpus callosum in some biopsies) and use the proportion of inhibitory cells as a technical factor we control for in our models.

We used the mixed-effects differential expression analysis framework DREAM^40^ to infer differentially expressed (DE) genes (adjusted p-value < 0.05) due to striatal region and/or learning, while accounting for random and fixed effects in our data. Important covariates such as hemisphere were included as fixed effects, and technical factors such as the proportion of inhibitory cells or library batch were included as random effects. To account for the non-independence of biopsies made within the same animal as well as the experimental design of learning mice and yoked controls, we also defined mouse pair as a random effects predictor (see Materials and Methods for model details). The dominating factor that explains variation in this dataset is striatal region (Figure 2D-E), likely reflecting known differences in the composition of cell types (e.g., D1 and D2) and the influence of spatial gradients^26,50^. This leads to approximately 13,000 DE genes among regions (i.e., ∼77 % of all detected genes), which is expected given the large statistical power to detect DE genes with 66 mice, each measured in both hemispheres.

Across the three striatal regions, the numbers of DE genes vary substantially by region: 10,941 in the VMS (5,179 upregulated, 5,762 downregulated relative to other regions), 10,431 in the DLS (5,486 upregulated, 4,945 downregulated), and 4,604 in the DMS (2,456 upregulated, 2,148 downregulated) (Extended Data Figure S5). Notably, the DMS shows the fewest DE genes, approximately half of those observed in either the VMS or DLS, positioning it as a transcriptionally intermediate region. This is consistent with a functional gradient spanning the ventromedial-to-dorsolateral axis. All region-wise findings can be explored in our interactive web application, but note that recent large-scale single-cell and spatial transcriptomics approaches are better suited for questions that require higher cellular and spatial resolution (see Discussion). This said, a few examples are worth highlighting as they illustrate well-established biology and validate our approach. *Pvalb*, a marker for the Parvalbumin-positive (PV+) fast-spiking interneurons is 3.6-fold higher expressed in the DLS than in the other regions (95% CI=[3.30, 3.93], adjusted p-value <10^-88^) as expected from previous studies^51,52^. Similarly, *Shank3*, a gene known to be implicated in the development of the Autism Spectrum Disorder^53,54^ is 1.09-fold higher expressed in the DLS than in the other regions (95% CI=[1.07, 1.11], adjusted p-value <10^-12^), which could be relevant when studying its function. Hence, we think the differences among regions that can be identified in our dataset, can be helpful to contextualise genes of interest within the striatal transcriptional landscape. However, the central aim and primary strength of this study is to probe learning-related transcriptional changes using a large number of biological replicates within a balanced experimental design. This approach enables us to quantitatively compare learning-related expression effects across regions and learning stages. For each comparison of interest, we apply the same analytical framework, which we outline before presenting the corresponding results.

### Analysis framework and the app

Using the mixed effects model, we analyse the following comparisons: “Striatal region” (regional differences within each mouse with all other factors included in the regression model), “Yoked stages” (comparison between yoked controls from the different stages), “Learning stages” (learners vs. yoked controls), “Between stages” (yoked-subtracted learning stage comparisons), “Learning effects VMS vs. DMS”, “Learning effects VMS vs. DLS” and “Learning effects DMS vs. DLS” (pairwise interaction between regions and learning effects). For each comparison, we quantify global expression shifts (mean absolute log-fold change), cluster samples based on their expression profiles, identify differentially expressed (DE) genes, identify DE-enriched gene sets, and identify changes in transcription factor (TF) activity. These analyses can be used to generate and follow-up hypotheses in various contexts and our interactive web application allows the user to adapt, filter, and download the requested data for all different comparisons we perform (Extended Data Figure S6).

Before highlighting interesting aspects of these analyses below, we think it is helpful to mention some general aspects for their interpretation: Our balanced design, with the same number of replicates across groups of learners and a similar amount of measurement noise expected across the striatal regions allows to assume a similar amount of statistical power^55^. This allows comparing the magnitude of expression changes by the number of DE genes or the average fold-change across conditions, e.g., asking whether more expression changes occur early or late during learning or whether more learning-dependent expression changes occur in the DMS or DLS. This could be regarded as analogous to comparing changes in electrical activity measured with microelectrodes or changes in oxygen saturation in an fMRI. This is a new way to interpret expression measures beyond the usual hypothesis generation and is possible only with large, balanced datasets like this one.

To describe DE genes for a given comparison on a more integrated level, we do a gene set enrichment analysis that uses gene annotations for biological processes provided by the Gene Ontology consortium^42,56^ and cell-type signatures and curated gene sets provided by the MSigDB collections^44^. Such Gene Set Enrichment Analysis (GSEA) tests whether a set of detected genes that share an annotation, has a higher number of DE genes than expected given all annotated DE and all annotated detected genes. We display the five most significantly enriched sets to avoid cherry picking. It is noteworthy that the top five in this list can change with changes in annotations of the reference databases or by using different DE thresholds, and that many gene functions are unknown and/or not annotated. This said, we think that it is still the best and most widely used possibility to describe the biology underlying DE genes.

Finally, we aim to gain insight into the transcription factors (TFs) that may drive the observed expression changes in a given comparison by estimating their activity. To this end, we use known regulatory relationships between TFs and their target genes from CollecTRI^45^ and assess whether the expression of these targets is consistent with the expected regulatory effects using decoupleR^46^. In brief, if a TF is known to activate and/or repress a set of target genes, its inferred activity is increased when the activated targets are more frequently up and/or downregulated than expected in a given condition. Conversely, its activity is inferred to be decreased when its targets change in the opposite manner. To increase the precision, one can filter the identified TFs for those that show a significant co-expression with their targets in our data.

In summary, we provide a robust and standardised analysis of expression changes in the learning striatum, designed to gauge overall magnitudes of expression change and to support hypothesis generation via an interactive, easy-to-use, open-access database.

### Mild gene expression changes in the control mice across time

We first investigated whether the time spent in isolated chambers induces gene expression changes. To this end, we compared the three yoked control groups, which did not perform the learning task, but were located in the same experimental room, solitarily housed in a control chamber and were fed the same amount of food as the learning mice (Extended Data Figure S7A). We hypothesised that if social isolation or other housing-related variables had a cumulative effect on striatal gene expression, we would expect the number of DE genes to increase with time spent in the chambers. However, we found only mild changes including 6 differently expressed (DE) genes when comparing yoked-intermediate against yoked-early mice, with a 95% percentile interval of 4.9-24.2 DE genes as assessed by jackknife resampling, and 0 [0.0-1.0] DE genes when comparing yoked-intermediate against yoked-late (Extended Data Figure S7B). These small transcriptional changes were also reflected in low average absolute log-fold changes (|LFC|) of 0.08 (y-Intermediate vs y-Early), 0.07 (y-Late vs y-Early) and 0.07 (y-Late vs y-Intermediate) (Extended Data Figure S7C). The six DE genes (Extended Data Figure S7E) are not commonly associated with canonical stress-response pathways and are not enriched in particular annotations (all GSEA test p-values>0.01). Hence, gene expression changes little over time in the control mice.

### The striatum shows a strong transcriptional response during early learning

As described above, we sampled 11 mice per learning stage, and each learning mouse was paired with a yoked control, i.e., an age- and sex-matched mouse, which was housed in a control chamber, received the same amount of food and was sacrificed in parallel (Figure 3A). Thus, apart from performing and learning the task, the learning mouse and its yoked control are as similar as we could possibly make them. Importantly, we also make use of this pairing in our statistical model (see Materials and Methods). We first compared each learning group (early, intermediate, and late) separately and accounted for region effects in our model to analyse the average effect across the three regions. We found a strong transcriptional response during the early phase of learning as early learners differed from their yoked-controls by 691 DE genes (95% percentile interval: [484.7, 701.5]). At intermediate and late learning stages, this difference dropped to 114 [47.0, 164.5] and 87 [40.0, 164.5] DE genes, respectively (Figure 3B). This pattern of a strong change in expression levels during early learning compared to intermediate and late learning is also seen by declining average expression changes (Figure 3C) and in a clustering pattern that tends to separate early learners from yoked controls (Extended Data Figure S8B). Notably, learning mice at the intermediate and late stage do not differ much, as no DE genes are found when comparing intermediate and late learning mice (after subtracting the respective yoked controls, see Methods and Extended Data Figure S9), while these two differ by 120 [79.0, 144.1] and 95 [61.9, 128.7] DE genes to the early stage, respectively.

**Figure 3.**
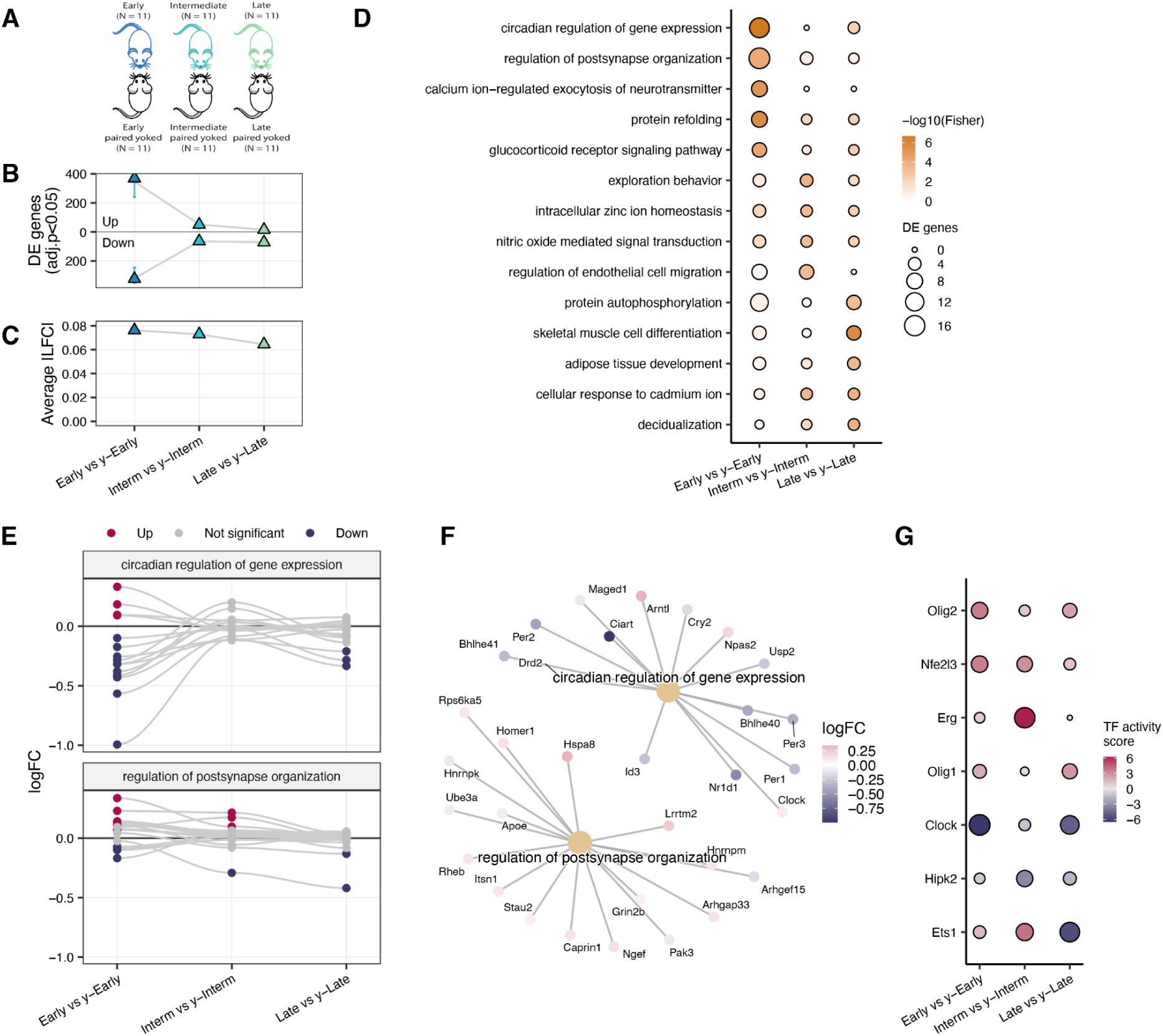
Learning-specific transcriptional responses in the striatum. A – Schematic overview of the experimental setup regarding learning mice across time and the yoked paired mice. B – The number of detected DE genes at each learning stage between learners and their yoked mice counterparts. Total number of DE genes: Early vs y-Early = 691; Intermediate vs y-Intermediate = 114; Late vs y-Late = 87. C – The average effect sizes (|LFC|) across all genes at each learning stage between learners and yoked mice. B,C -Sampling distribution was generated using jackknifing, e.g., each mouse pair was removed once and the same model was refitted. Error bars: 95% percentile intervals. D – Top five most significant gene ontology categories per learning stage and their enrichment in the other stages (Biological process, Fisher’s p-value<0.01). E – Expression differences of significant genes between learning and yoked mice from selected GO categories. F – Genes in pathways from E in the comparison between Early learners and their yoked counterparts. The nodes represent GO categories and the edges represent significantly DE genes belonging to the respective categories, colored according to their log-fold change. G – Predicted transcription factor (TF) activity differences between the respective learning stage and yoked mice groups using the CollecTRI database. These associations were further pruned by the co-expression between the TF and the targets compared to TF and random genes (one-sided Wilcoxon rank sum test p-value < 0.05).

These DE genes underlie the enrichment of 73, 23 and 28 GO categories at the early, intermediate and late stages, respectively. Some of the GO categories most enriched for DE genes in early learning include *circadian regulation of gene expression*, *regulation of postsynapse organization*, and *calcium ion-regulated exocytosis of neurotransmitters* (the most enriched categories are illustrated in Figure 3D, and the full list is available in the web application). While one must be careful not to overinterpret such term listings (see the section on the analysis framework above), it is fair to say that terms like postsynapse organisation or exocytosis of neurotransmitter confirm expectations about gene expression changes during (early) learning whereas circadian rhythm regulation might be a more unexpected finding. But note that categories can share substantial gene-level overlap. For example, *Drd1*, *Drd2*, and *Adora2a,* which are central to dopaminergic signalling, and, hence, also confirm expectations for striatal learning, are DE genes annotated in *circadian behavior, regulation of synaptic plasticity, or locomotory behaviour*, to name a few. To illustrate such relations in more detail, we show which genes they contain and how they are differentially expressed during learning (examples in Figure 3E-F, Extended Data Figure S6). This said, it is worth pointing out that circadian mechanisms do play a critical role in learning and cognitive processes^57,58^ and a central hub of this regulation *Arntl* (also known as *Bmal1*) is an upregulated DE gene during early learning in our data and has recently been shown to directly influence striatal dopamine signalling^59^. Circadian regulation pathways are also among the most enriched gene sets when using the M2 annotations from MSigDB (Extended Data Figure S8C) that comprise gene sets of pathways curated from the literature^44^. Notably, these pathways are mostly not enriched at the intermediate and late stages of learning. At these stages, learning related categories like *adipose tissue development* are enriched (Figure 3D), but also terms like *long-term synaptic potentiation, vasculogenesis*, and *regulation of endothelial cell migration* emerge and could indicate slower structural or metabolic adjustments that continue after behavioural performance has stabilized.

Another way to explore the expression changes, is to identify transcription factors that have increased or decreased activity by testing whether the expression of their known targets changes accordingly. When also filtering for significant correlations of the TF and its targets in our data, we identify seven TFs with changed activity in at least one of the three learning stages (Figure 3G). One notable finding is the increased activity of *Olig1* and *Olig2* that are core regulators of oligodendrocyte lineage specification^60^. Their increased activity may reflect the fact that neuronal activity and learning are known to promote oligodendrocyte differentiation and adaptive myelination^61,62^. Another finding is the increased activity of *Ets1* and *Erg* at the intermediate learning stage. Both transcription factors are known regulators of vascular development and angiogenesis^63,64^. Their increased activity may therefore reflect enhanced vascular remodeling or blood vessel formation, a process that has been shown to increase in response to neuronal activity in the sensory cortex^65^. Clearly, more could be said about the hundreds of DE genes and their biological associations, but we hope that these examples illustrate how the data can be explored and interpreted to corroborate existing hypotheses and generate new ones.

### Learning-induced transcriptional changes are largely shared across striatal regions

In the section above, we have only analysed learning signatures that are common across the regions. To test whether there is evidence for learning signatures that are region-specific, we added a learning-region interaction term to our model (Model 2, see also Materials and Methods). We did not find any genes that showed a significant region-specific learning effect (all adjusted p-values > 0.05). This is remarkable given the large differences between regions (Figure 2E and Extended Data Figure S5) and the fairly strong effects of learning (Figure 3). However, we also have more statistical power to detect those latter changes as we effectively combine more samples per group.

To investigate whether the transcriptional response to learning is indeed consistent across regions when analysed independently, we split the expression matrices per region and compared each learning stage to its yoked control group (Model 3 in Methods). As expected, we found fewer DE genes than when combining all regions (138 versus 818) and almost all of them at the early learning stage (57 [35.0, 59.0], 48 [36.0, 60.0] and 74 [46.1, 89.0] for VMS, DMS and DLS, respectively; Figure 4A). Hence, neither the number of DE genes (Figure 4A) nor the effect sizes (Figure 4B) indicate region-specific differences across learning stages, including no evidence for a shift from DMS to DLS as learning transitions from goal-directed to habitual, as reported in neurophysiological studies^7,8^. While this does not exclude more subtle region-specific expression changes that may be detectable with increased statistical power, it indicates that most expression changes during striatal learning are region-independent.

**Figure 4.**
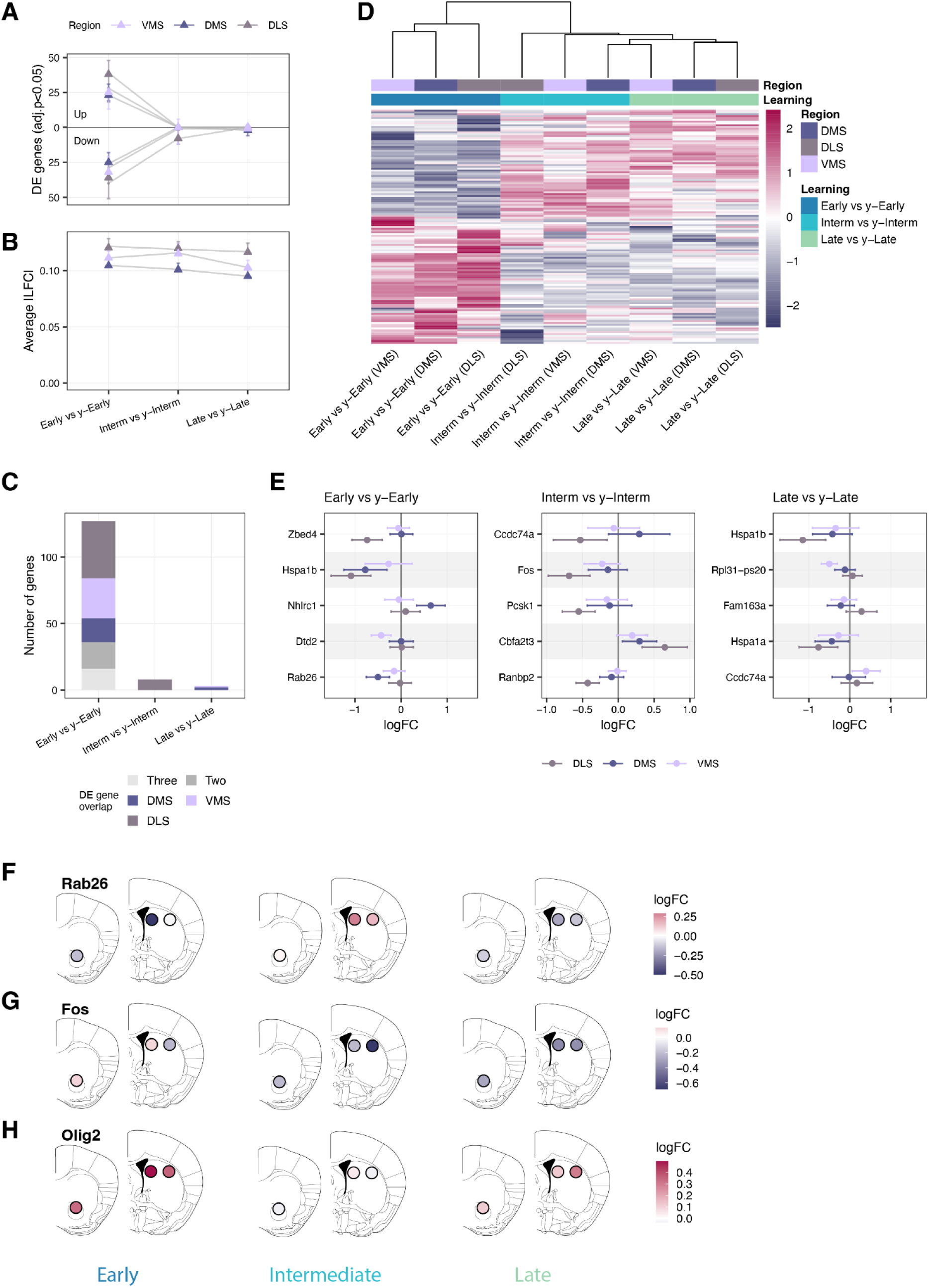
Learning-induced expression changes are not distinctively striatal region-specific. A – The number of detected DE genes at each learning stage between learners and their yoked mice counterparts for each region independently. On the level of gene expression, all striatal regions show most changes at the early learning stage. The total number of DE genes per region during the early stage is VMS = 57, DMS = 48 and DLS = 74. B – The average effect sizes (|LFC|) across all genes at each learning stage between learners and yoked mice per region. C – Number of DE genes per learning stage, stratified by regional overlap. Colours indicate genes detected exclusively in VMS, DMS, or DLS, or shared between two or all three regions. D – Clustering of learning genes (n = 138, adjusted p-value in any condition < 0.05) based on estimated logFC per learning stage per region. Distance = Euclidean, method = complete. Contrasts cluster primarily by the learning stage, not by region. E – LogFC estimates per region for the top 5 genes with the largest inter-regional variance in learning response, shown separately for each learning stage. Points represent estimated logFC; horizontal lines indicate 95% confidence intervals. F, G, H – Regional logFC maps for selected genes of interest across early, intermediate and late learning stages. Circle size and colour intensity reflect the magnitude and direction of the logFC relative to yoked controls (red: upregulated; blue: downregulated; white: no change). F – Rab26. G – Fos. H – Olig2. (VMS: AP = 1.10, ML = ±1.00, DV = 4.00; DMS: AP = 0.14, ML = ±1.50, DV = 2.10; DLS: AP = 0.14, ML = ±2.50, DV = 2.50).

To explore the most pronounced potential region-specific effects, we selected, at each stage, the 25 genes with the largest variation in log-fold change across striatal regions in response to learning. The top five most variable genes per learning stage are shown in Figure 4E, with an extended list provided in Extended Data Figure S10. To illustrate some of these region-specific dynamics, we highlight three genes of established relevance to learning and plasticity. For instance, *Rab26,* a Rab GTPase regulating the endocytosis and autophagic degradation of the serotonin transporter SERT^66^, showed a pronounced downregulation in the DMS and a milder reduction in the VMS. At the intermediate stage, this pattern reversed markedly in the DMS, which exhibited strong upregulation, with the DLS also showing modest upregulation, while the VMS returned to baseline. At the late stage, all three regions showed a mild downregulation (Figure 4F). Given that *Rab26* promotes SERT endocytosis, thereby reducing serotonin reuptake and increasing synaptic serotonin availability, this dynamic pattern raises the possibility of region- and stage-specific modulation of serotonergic signalling across the course of learning, with the DMS emerging as the primary site of regulation, consistent with the established role of DMS serotonin in encoding reward value and anticipation during associative learning^67^. On the other hand, *Fos*, a well-known immediate early gene reflecting neuronal activity^68,69^, showed mild upregulation in VMS and DMS during early learning, followed by slight downregulation at intermediate and late stages. In contrast, *Fos* expression in DLS was slightly downregulated during the early learning phase, dropped further at the intermediate stage, and remained moderately suppressed during late learning (Fig. 4G). Finally, *Olig2*, a regulator of oligodendrocyte differentiation^60^, exhibited a marked upregulation across all regions during early learning, with the DMS showing the strongest response. This was followed by a return to baseline at the intermediate stage, and a region-specific rebound at late learning, particularly pronounced in the DLS (Fig. 4F), suggesting potential glial engagement during consolidation. Despite limited statistical support at this point, these patterns suggest that our data capture potentially relevant moderate region-specific expression changes during striatal learning, which might be helpful in future investigations of striatal learning.

## Discussion

In the present study, we provide a robust molecular characterization of striatal learning in a well-controlled experimental design, enabled by combining a fully automated behavioural paradigm with efficient bulk RNA-sequencing of 396 striatal biopsies from 66 mice. This offers a detailed view on gene expression dynamics throughout early, intermediate, and late learning stages in three striatal regions. Crucially, each learning animal was paired with a yoked control matched for sex, age, genotype, and environmental conditions, allowing us to isolate transcriptional changes specifically attributable to learning. This represents, to our knowledge, the first temporal and spatial transcriptomic atlas of striatal learning at this scale, offering a valuable resource for exploring the molecular architecture of adaptive behaviour. We discuss the advantages and limitations of the methods and the resulting resource in turn.

### A new automated operant chamber

Operant conditioning set-ups have been used and developed since their creation in the 19^th^ century ^70,71^. Despite varying designs and complexity of operant learning paradigms, the vast majority of experimental work still relies on daily conditioning sessions of rather short duration. Such design often includes drastic experimental interferences to elevate the motivational drive of experimental animals to engage in the task such as, e.g., water- or food-restriction in mammals^72^. The automated home-cage behavioural paradigm used in our approach overcomes limitations of such traditional operant conditioning tasks. By allowing animals to live within the apparatus and engage with the task at their own pace and according to their own motivation, the system enables the collection of large numbers of trials over extended periods and provides insights into more naturalistic behaviour than demonstrated under classical, often food- or water-deprived operant conditioning paradigms. The large volume of data collected enables fine-grained analyses of learning from its early acquisition to later, overtrained stages, including the identification of alternative strategies that complement the ultimately most efficient and dominant action-outcome association. The high degree of automation furthermore standardises the task and ensures reproducible measurements across animals and experimental sessions, while enabling high-throughput data collection. This was essential for generating the large number of matched striatal samples (n = 396) required to extract learning-specific expression profiling signatures.

### A new framework for expression profiling

When measuring gene expression profiles, particularly in heterogeneous populations, single-cell resolution is generally preferred as cells represent fundamental, well-interpretable units of gene expression. Moreover, the spatial context of cells is often critical, making rapidly advancing spatial transcriptomic approaches highly attractive^73^. Combining cell-type annotations from genome-wide single-cell expression profiles with targeted spatial transcriptomics is an increasingly common strategy to characterize tissues, as exemplified by studies of GABAergic neurons in the mouse brain^26,50^. However, when comparing gene expression across conditions, as done here for learning, the signal-to-noise ratio is typically much lower than the differences between cell types and states captured in atlas-type studies. Consequently, robust conclusions require more biological replicates, rendering single-cell and, even more so, spatial transcriptomic studies cost-prohibitive for many laboratories. The exact costs depend strongly on the single-cell technology used and the extent to which biological replicates can be multiplexed^74^. However, taking 10x Genomics as a current standard, even the latest FLEX workflows, which enable substantially higher multiplexing, remain ∼100-fold more expensive per sample than the bulk RNA-seq approach used here. In addition, the generation of single-cell suspensions from adult brain tissue is not feasible, necessitating single-nucleus RNA sequencing and further increasing costs, sample preparation complexity, and susceptibility to technical variation and isolation biases compared to bulk biopsies. While one can expect that these aspects will improve in coming years, a study like done here would be currently technically substantially more challenging and cost-intensive when done with single-cell resolution. Furthermore, it is challenging but not impossible to infer the cellular origins of expression changes from bulk profiles. Some expression changes, such as those associated with oligodendrocyte maturation or vascularization, have clear candidate cell types. Furthermore, the growing availability of cell-type-specific reference data will increasingly enable deconvolution of bulk expression profiles, improving their interpretation^75^. Alternatively, bulk-derived changes can be followed up using hybridization-based spatial transcriptomics, as done after snRNA-seq studies^26,50^, although this is not yet well established for the smaller signal-to-noise ratios examined here.

Independent of whether bulk or single-nucleus approaches are used, a well-balanced experimental design with sufficient biological replication and proper randomization of factors of interest is essential. Both biological (e.g. circadian rhythm, biopsy location, age and sex) and technical factors (e.g. library batch, sequencing batch and processing time) can introduce effects comparable in magnitude to the signal of interest. A balanced experimental design with similar statistical power for factors of interest, also allows to interpret the magnitude of change e.g. by the number of differently expressed genes. This quantitative aspect of expression profiles can be viewed as a comparison of molecular activities, analogous in some respects to activity comparisons e.g. from fMRI signals.

In summary, the cost-efficiency of (bulk) RNA sequencing now enables the investigation of molecular processes across much larger sample sizes, facilitating the detection of subtle effects and enabling quantitative comparisons across conditions when combined in a larger, well-balanced experimental design. We think that such a framework of expression profiling will be helpful for many biological questions that profit from extending beyond two-way comparisons.

### Magnitudes of expression changes during striatal learning

Combining the automated operant chamber and efficient RNA-seq profiling, we investigated how three subregions in the striatum change expression profile over three learning stages in a visual discrimination task. Overall, we find a relatively strong transcriptional response (691 DE genes) during early acquisition that is reduced at intermediate (120) and late (95) learning stages that differ little from each other (0). These responses are very similar across the three striatal subregions and we find no genes that show a significant region-learning interaction. This latter finding is particularly intriguing in light of prior neurophysiological studies, which have long suggested that striatal regions are differentially recruited across learning phases^7,8^. One possibility to reconcile these findings is that differential recruitment of regions at the neurophysiological level arises from pre-existing molecular differences, for example established during development. These differences shape different neuronal circuitries, leading to differential electrophysiological recruitment on a background of otherwise similar (expression) activity across regions. More work will be needed to disentangle how the magnitude of expression change is connected to the magnitude of neuronal activity and how this can be linked to differential recruitment of striatal regions at different learning phases.

### Composition of expression changes during striatal learning

The composition of expression changes is highly informative, as many genes and gene sets are associated at least to some extent with interpretable biological processes. Given the known complexity and diversity of these processes during striatal learning, we found it appropriate to present the data as a resource that can be explored interactively. That said, we highlight several patterns that serve two purposes: first, to contextualize the data within current models of striatal function, and second, to illustrate the types of integrative analyses enabled by this resource. We concentrate the discussion on three findings that we consider particularly informative, as they extend beyond neuron-centric models of learning and highlight coordinated contributions from glial, vascular, and temporal regulatory processes.

One of the most striking observations emerging from the atlas is the robust enrichment of circadian-related gene sets during early learning, supported by both GO enrichment^42,56^ and curated MSigDB analyses^44^. Because functional categories such as “circadian rhythm” can include heterogeneous genes, including dopaminergic (Fig. 3F), their interpretation benefits from gene-level inspection. The atlas is specifically designed to enable this type of interrogation. For instance, inspection of the canonical *Clock* gene^76^ indicates stage-specific up-regulation during early learning, with no effects maintained at intermediate or late stages. Circadian processes have previously been linked to learning consolidation^58^, cognitive evolution^57^, and psychiatric vulnerability^77,78^. Together, these observations suggest that early associative learning engages central circadian transcriptional programs, potentially aligning neuronal plasticity with temporal regulatory mechanisms.

In contrast, vascular-related gene sets emerged during intermediate and late learning stages. GO enrichment analyses^42,56^ indicated increased representation of vasculogenesis-associated transcripts during these later phases. While vascular changes are often interpreted as secondary metabolic correlates of neural activity, recent work has highlighted potential neuromodulatory roles of hemodynamics itself, through mechanical, thermal, or diffusible blood-borne factors^79^. Consistent with this view, learning has been shown to require angiogenesis but not neurogenesis^80^, supporting a late-stage vascular contribution to consolidation. Within the temporal framework observed here, these findings suggest that vascular remodeling may represent a delayed structural adaptation that accompanies circuit stabilization rather than an immediate correlate of early synaptic plasticity.

A third prominent feature of the dataset concerns oligodendrocyte lineage–associated transcription factors. At the global striatal level, *Nfe2l3*, *Olig1*, and *Olig2* showed elevated activity during early learning. Notably, *Olig1* and *Olig2* activity returned to near-baseline levels at the intermediate stage before re-emerging at late learning. Regionally resolved analyses revealed further nuance: *Olig2* was upregulated across all striatal subregions during early learning, with particularly strong effects in the DMS. At late stages, a rebound was especially pronounced in the DLS, a region classically implicated in automatization^8,9^. These dynamics are consistent with emerging evidence linking oligodendrocyte lineage cells and adaptive myelination to learning consolidation ^61,62^. The biphasic pattern observed here may reflect an early priming phase, followed by a transient restraint, and later structural reinforcement as behaviour becomes stabilized. Although this interpretation remains speculative, the temporal and regional specificity of the signal suggests that glial plasticity is not merely a passive consequence of neuronal activity but may participate in orchestrating long-term circuit optimization.

Taken together, these three observations point toward a temporally structured and multi-compartment response during striatal learning. Early stages are marked by engagement of circadian regulatory programs alongside activity-dependent transcription, potentially coordinating neuronal plasticity with broader temporal dynamics. Intermediate and late stages instead show signatures of vascular remodeling, consistent with structural and metabolic adaptation accompanying consolidation. Finally, the biphasic engagement of oligodendrocyte lineage programs suggests progressive glial contributions to long-term circuit refinement. Rather than reflecting isolated neuronal signalling changes, the transcriptional landscape uncovered here supports a model in which learning emerges from coordinated interactions.

### Limitations and outlook

While our dataset represents a substantial resource, it is currently limited in its spatial resolution as discussed above. But also increasing its temporal resolution, especially during early learning phases, should be informative. Another axis is to better link gene expression during learning to upstream phenotypic modalities, such as chromatin states and downstream phenotypic modalities, such as protein levels and neurophysiological properties. Finally, further work is needed to assess how these patterns generalize across different tasks and control conditions.

Despite these limitations, the overarching structure of our findings remains clear. The combination of strong baseline regionalisation with a conserved temporal learning trajectory provides a unifying explanatory model. It suggests that long-standing physiological specialisations of the ventral-to-dorsal striatum arise not from discrete molecular programmes during learning but from region-specific baseline identities operating within a shared temporal transcriptional cascade. This framework also situates neuropsychiatric risk genes, such as *Shank3*, for example, within defined striatal regions, offering mechanistic avenues for interpreting their contributions to behavioural vulnerability. The interactive atlas accompanying this work is intended as a community resource for querying genes, pathways, and regulators across regions and learning stages, and for facilitating cross-study integration. In sum, our results reconcile classical functional specialisation with a unified temporal program of striatal learning, identifying when, and not only where, transcriptional plasticity is deployed during the transition from acquisition to consolidation.

## Supporting information

Extended Data Table 1 GLMM

Extended Data Table 2 GLM

## Data Availability

Raw RNA-seq data generated in this study is available at: https://www.ebi.ac.uk/biostudies/arrayexpress/studies/E-MTAB-15694

Behavioural data and analysis scripts are available at: https://github.com/elianalousada/Expression_profiling_of_the_learning_striatum

Expression data and analysis scripts are available at: https://github.com/zkliesmete/Dopaloops

Web application: https://shiny.bio.lmu.de/Dopaloops/

## Contributions

CS, WE, and EBu conceptualised the research; EL conducted experiments and analysed behavioural data; AJ, EBr, DR, LEW generated and preprocessed expression data; ZK, IH analysed expression data; EL, ZK, CS, WE wrote the manuscript.

## Acknowledgements

All animal work was conducted at the PHENO-ICMice facility. This work furthermore benefitted from the equipment and services from the iGenSeq core facility at ICM for the genotyping of the animals. This work was realised with the following funding: Agence Nationale de la Recherche (ANR-19-ICM-DOPALOOPS) (EBu, WE), the Fondation pour la Recherche Medicale (FDT202204015143) (EL), and the L’Oréal-UNESCO Fellowship 2016 (CS). The core facilities were supported by “Investissements d’avenir” (ANR-10-IAIHU-06 and ANR-11-INBS-0011-NeurATRIS) and “Fondation pour la Recherche Médicale”. We also thank Leonhard Schaffmayer for helping with the neurobiological interpretation of the results.

## Extended Data: Figures with legends

**Extended Data Figure S1.**
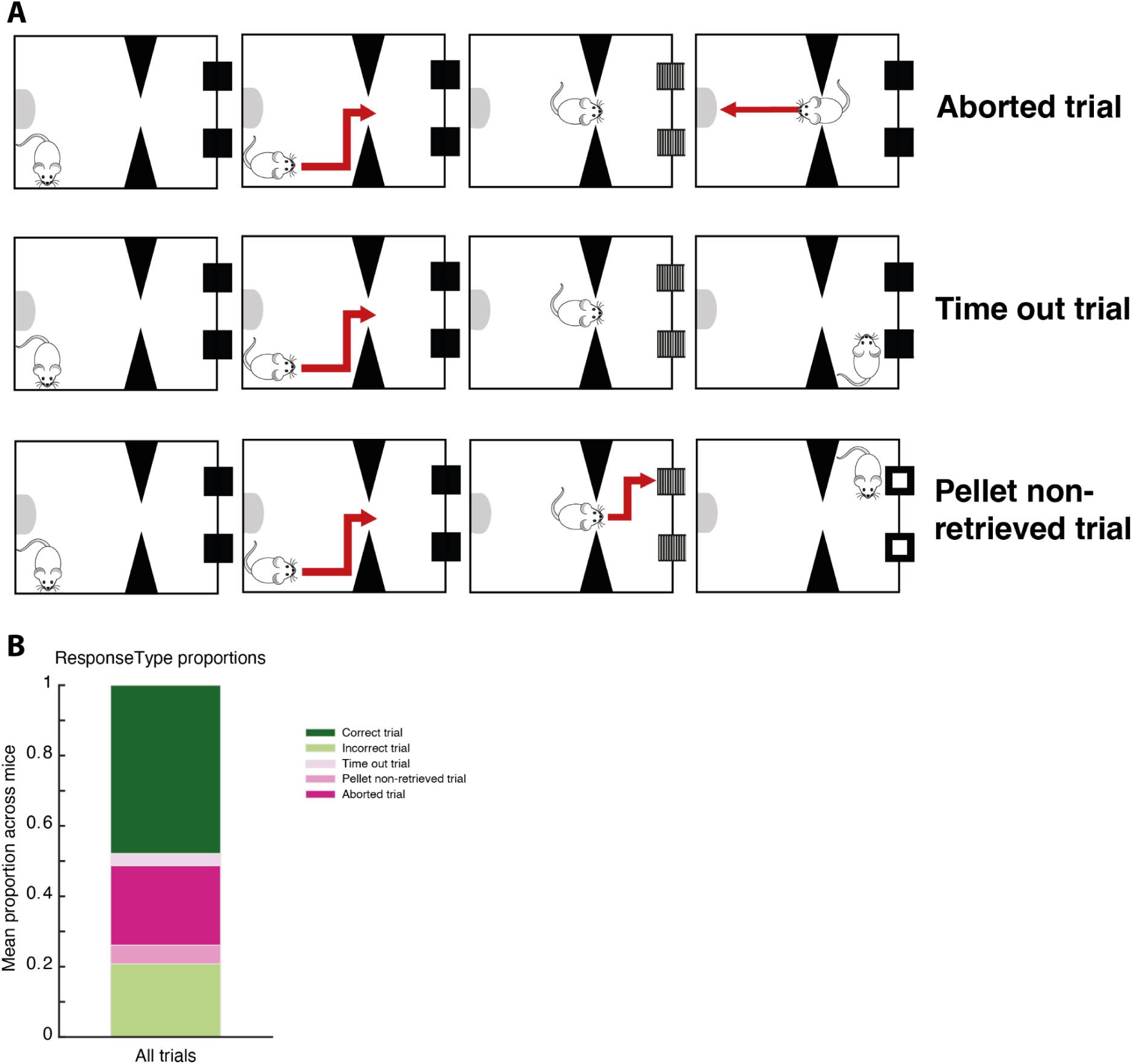
Behavioural task details. A – Trial types. In addition to correct and incorrect responses, three other trial categories were defined: *aborted trials* (entry and exit from screen compartment within 60 s without response), *time-out trials* (no response within 60 s), and *pellet non-retrieved trials* (reward not retrieved within 15 s of a correct response). B – Proportion of trial types averaged across mice, during the entire task.

**Extended Data Figure S2.**
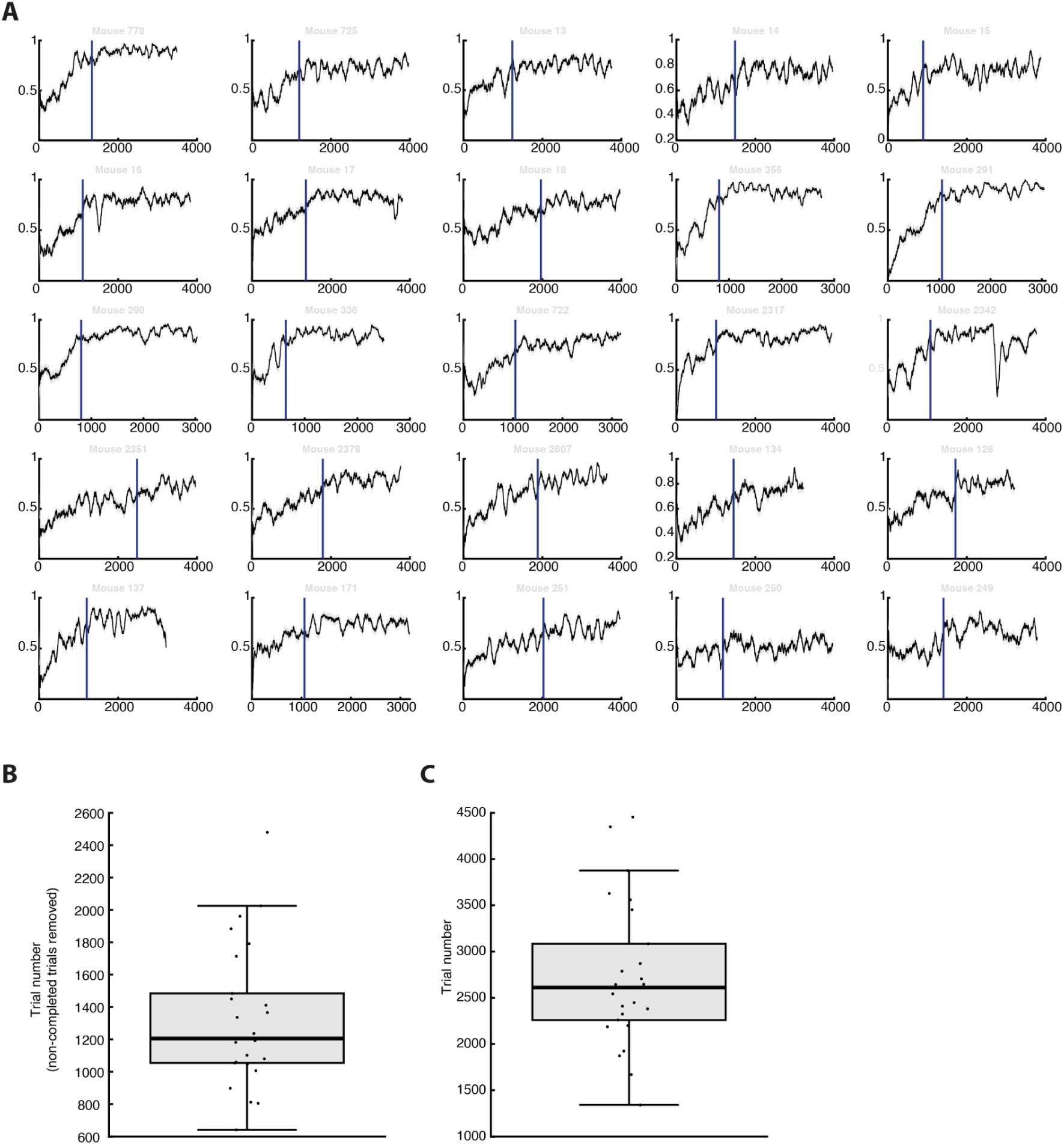
Definition of the time point of the early learning stage. A – Performance curves for each mouse (n = 25) across trials. Performance was computed as a sliding-window average of correct responses. The vertical blue line indicates the onset of stable performance (plateau), defined as the first sustained block of trials meeting predefined stability criteria (see Methods). This time point was used to determine the individual transition to the early learning stage. B – Distribution of early learning time points expressed as trial number after excluding non-completed trials (i.e., including only correct and incorrect responses). Boxplots show the median (center line), interquartile range (box), and full range (whiskers), with individual mice overlaid as dots. C – Distribution of early learning time points expressed as absolute trial number (including all trial types). Boxplots are displayed as in panel B.

**Extended Data Figure S3.**
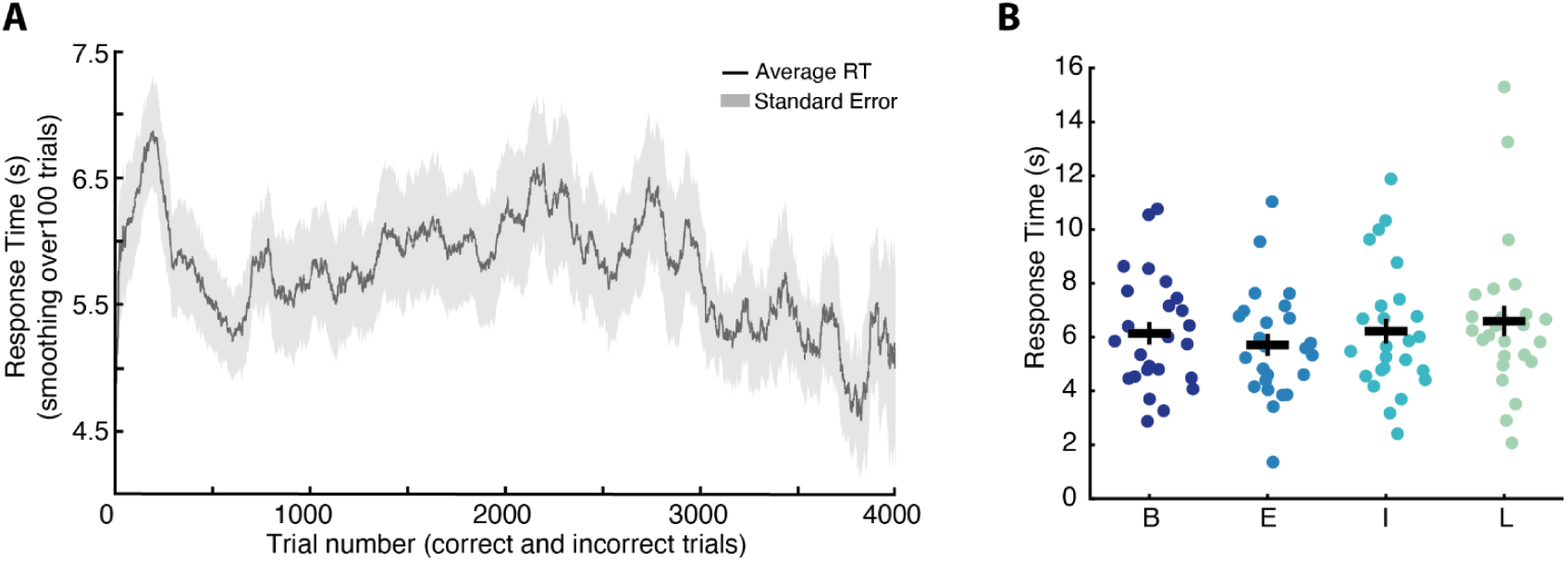
No learning-stage related difference in response times. A – Global response time (RT), defined as the interval between stimulus onset and touchscreen response, is highly variable across animals and does not systematically differ across learning stages. Shown is mean ± SEM across trials. B – Individual-level data across learning stages for RT. No significant differences between stages.

**Extended Data Figure S4.**
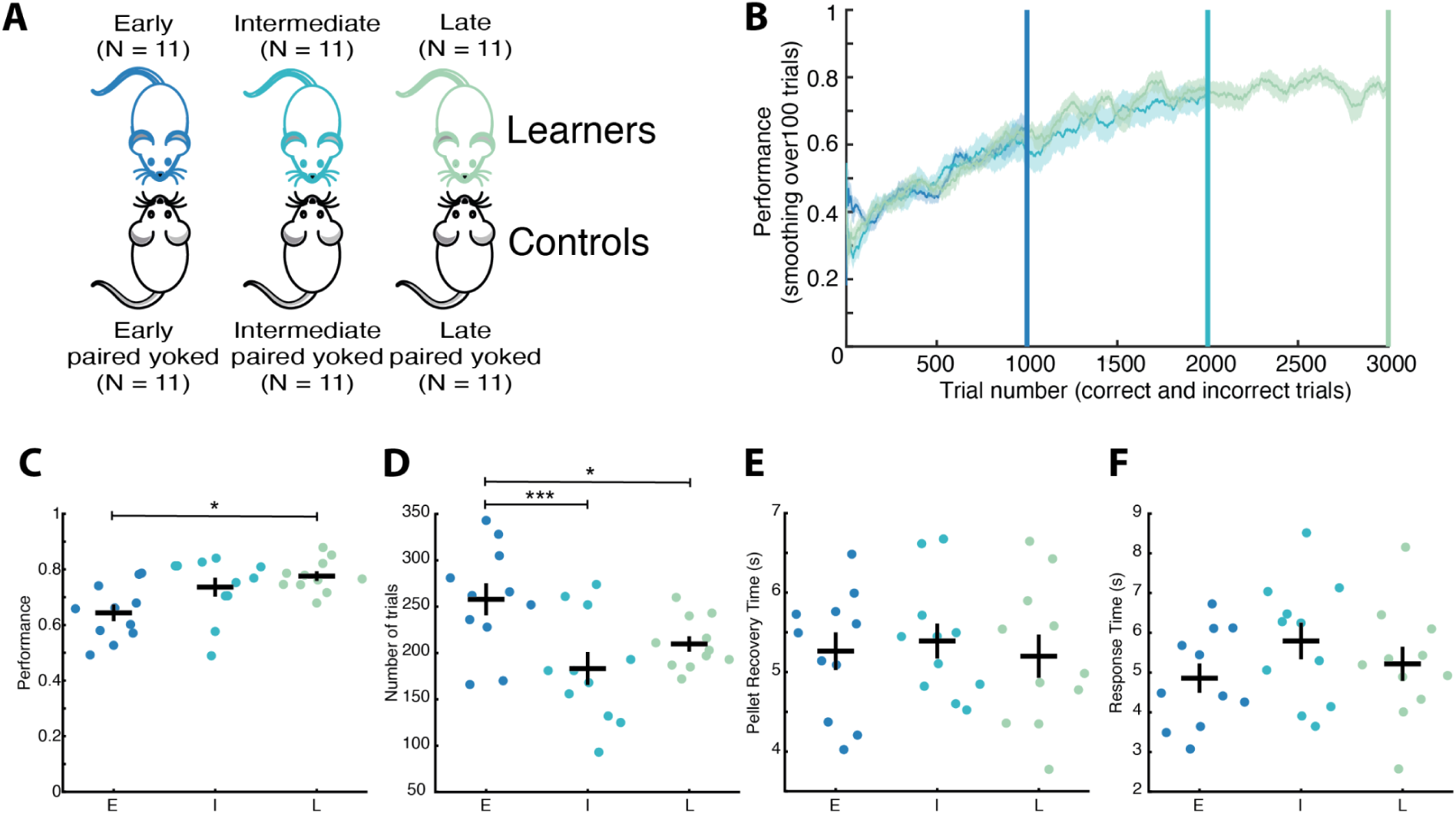
Replication of behavioural readout parameters in animals applied for molecular mapping. A – Illustration of experimental design. Learners were paired with yoked controls to account for non-task-related variables. B – Mean performance (± SEM) of each learning stage group up to the respective stage criterion: 1000, 2000, and 3000 completed trials for early, intermediate, and late learners, respectively. Coloured lines represent the three defined learning stages: early (blue), intermediate (turquoise), and late (green). C – Individual data points of performance. Significant increase between early and late learning stages. D – Individual data points of number of initiated trials. Significant decrease between early and intermediate, and late learners. E, F – Individual data points of PRT and RT, respectively. Absence of significant differences across all pairwise comparisons. Early, intermediate and late learners are colour coded in blue, turquoise, and green, respectively. E: early, I: intermediate, L: late. Statistical significance levels: p < 0.001 (***), p < 0.01 (**), p < 0.05 (*).

**Extended Data Figure S5.**
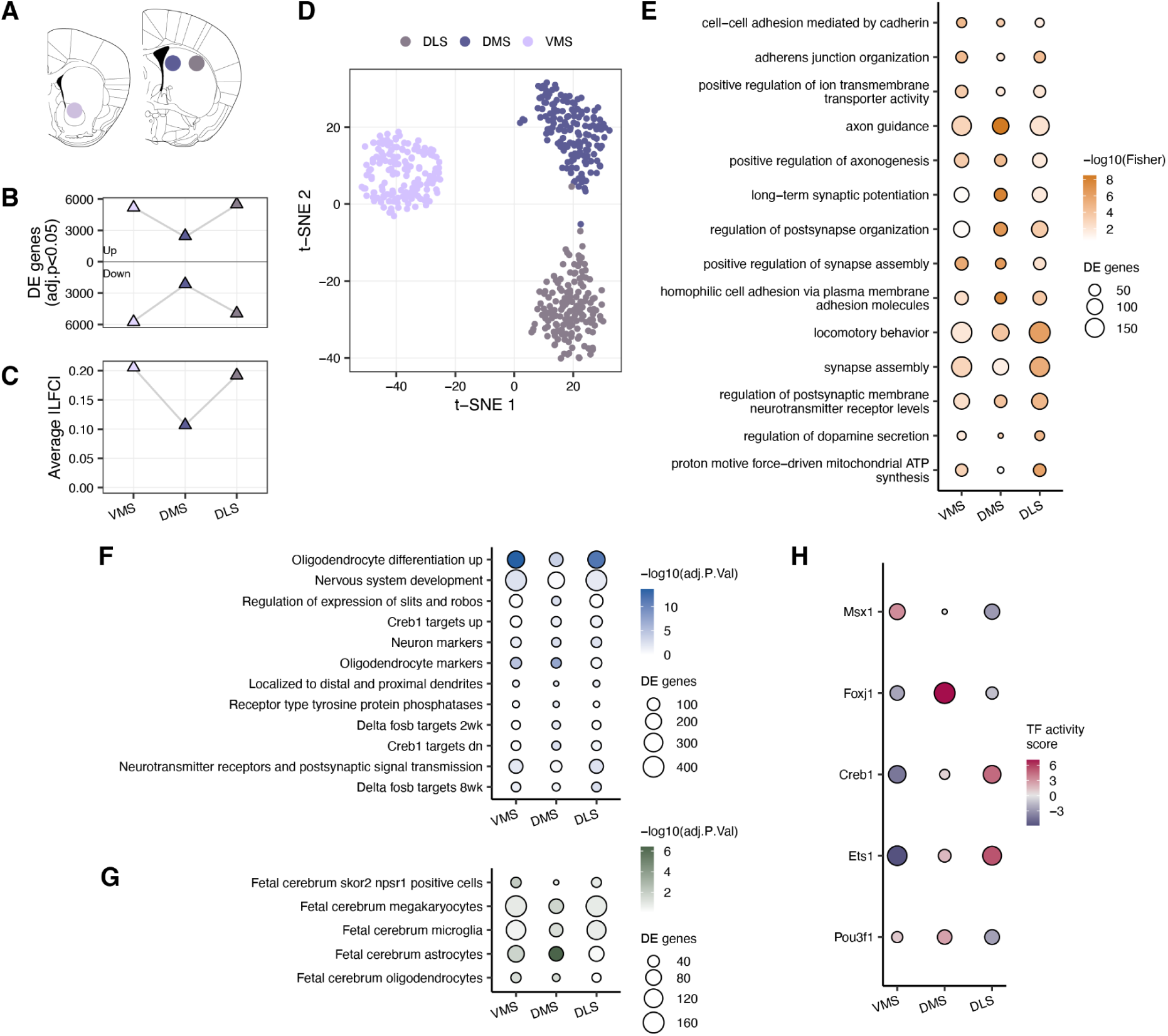
Expression differences between striatal regions. A – Schematic overview of the three striatal regions included in the study (VMS: AP = 1.10, ML = ±1.00, DV = 4.00; DMS: AP = 0.14, ML = ±1.50, DV = 2.10; DLS: AP = 0.14, ML = ±2.50, DV = 2.50). VMS is depicted in light purple, DMS in dark blue, DLS in grey. B – The number of detected DE genes between each striatal region compared to the other two. Total number of DE genes: VMS vs others = 10,941; DMS vs others = 4,604; DLS vs others = 10,431. C – The average effect sizes (|LFC|) across all genes between each striatal region and the other two. B,C –Sampling distribution was generated using jackknifing, e.g., each mouse pair was removed once and the same model was refitted. Error bars: 95% percentile interval. D – Dimensionality reduction using t-SNE based on the 1000 most significant DE genes (PCs = 50, perplexity = 20, theta = 0.1). All random effect predictors (see methods) have been regressed out prior to clustering. DMS is least distinct from the other striatal regions, consistent with the observations in B,C. E – Top five most significant gene ontology categories per region and their enrichment in the other regions (Biological process, Fisher’s p-value<0.01). F – Enriched M2 categories from selected published annotations on function in the mouse, between the different striatal regions (BH-adjusted p-value<0.05). G – Enriched M8 categories from selected published annotations on cell type markers in the mouse, between the different striatal regions (BH-adjusted p-value<0.05). H – Predicted transcription factor (TF) activity differences between the respective striatal region and the others using the CollecTRI database. These associations were further pruned by the co-expression between the TF and the targets compared to TF and random genes (one-sided Wilcoxon rank sum test p-value < 0.05).

**Extended Data Figure S6.**
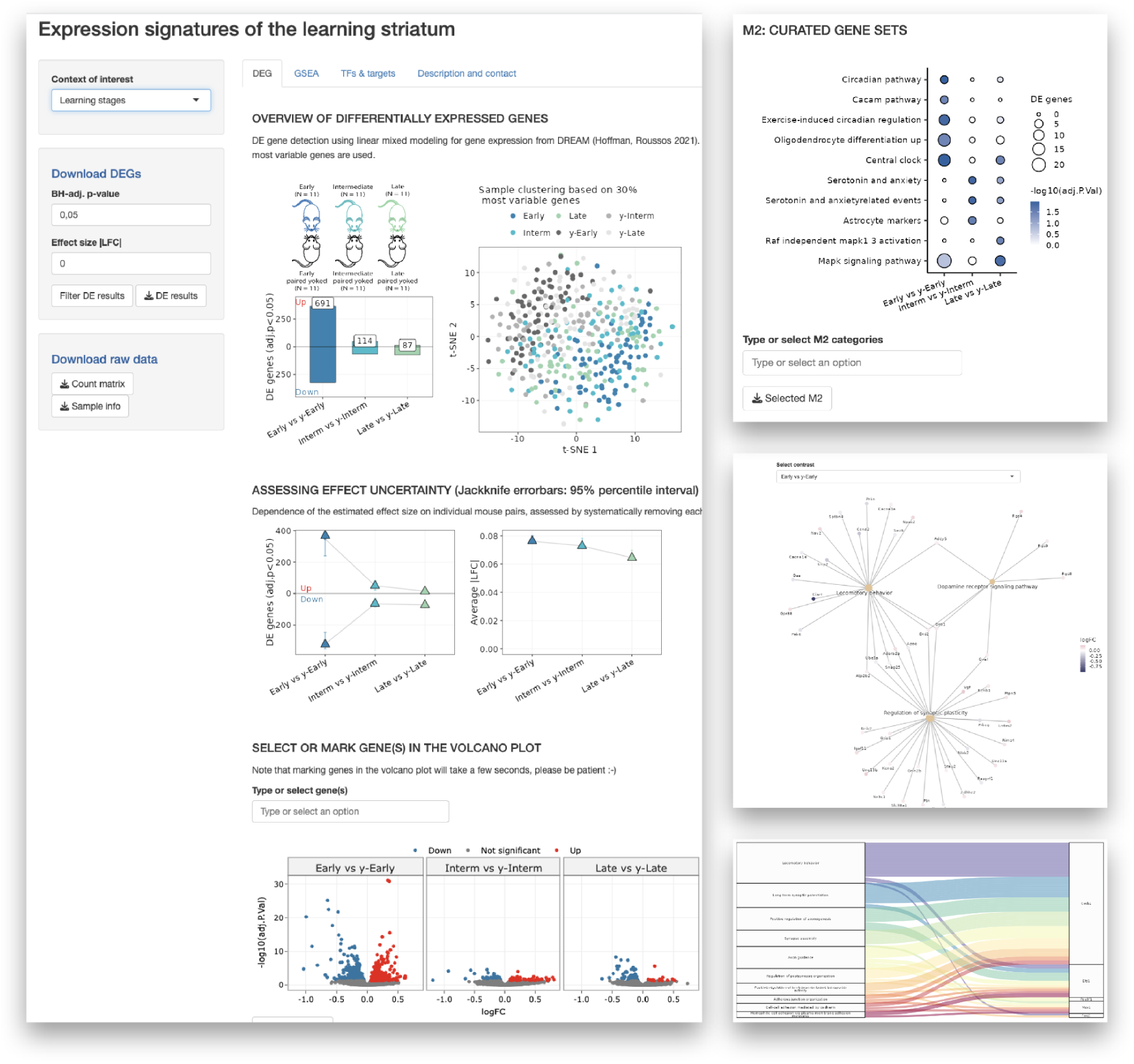
Interactive web application for exploring the expression atlas of the learning striatum. The application is publicly accessible at https://shiny.bio.lmu.de/Dopaloops/ and allows users to interactively query, visualise, and download all results generated in this study. The left panel allows selection of the contrast of interest (here: learning stages) and provides options to filter results and download the full list of DE genes for independent downstream analysis. The main panel is organised into four tabs: DEGs (differentially expressed genes), GSEA (gene set enrichment analysis), TFs & Targets (transcription factor activity), and Description & Contact (study details and correspondence information). The DEG tab is shown here and includes: the number of up-and downregulated DE genes per learning stage; a t-SNE dimensionality reduction plot of expression profiles; logFC across learning stages; and volcano plots for each contrast, in which individual genes of interest can be selected to visualise their logFC and significance across all learning stages simultaneously. The top right panel displays enrichment results for M2 curated gene sets from MSigDB. Below it, a genes in pathway plot illustrates the genes significantly enriched in GO categories. The bottom right panel depicts a Sankey plot showing the overlap between GO categories (left) and TF activities (right) in terms of the underlying enriched DE genes that are annotated as specific TF targets. This and more can be visualized using the app.

**Extended Data Figure S7.**
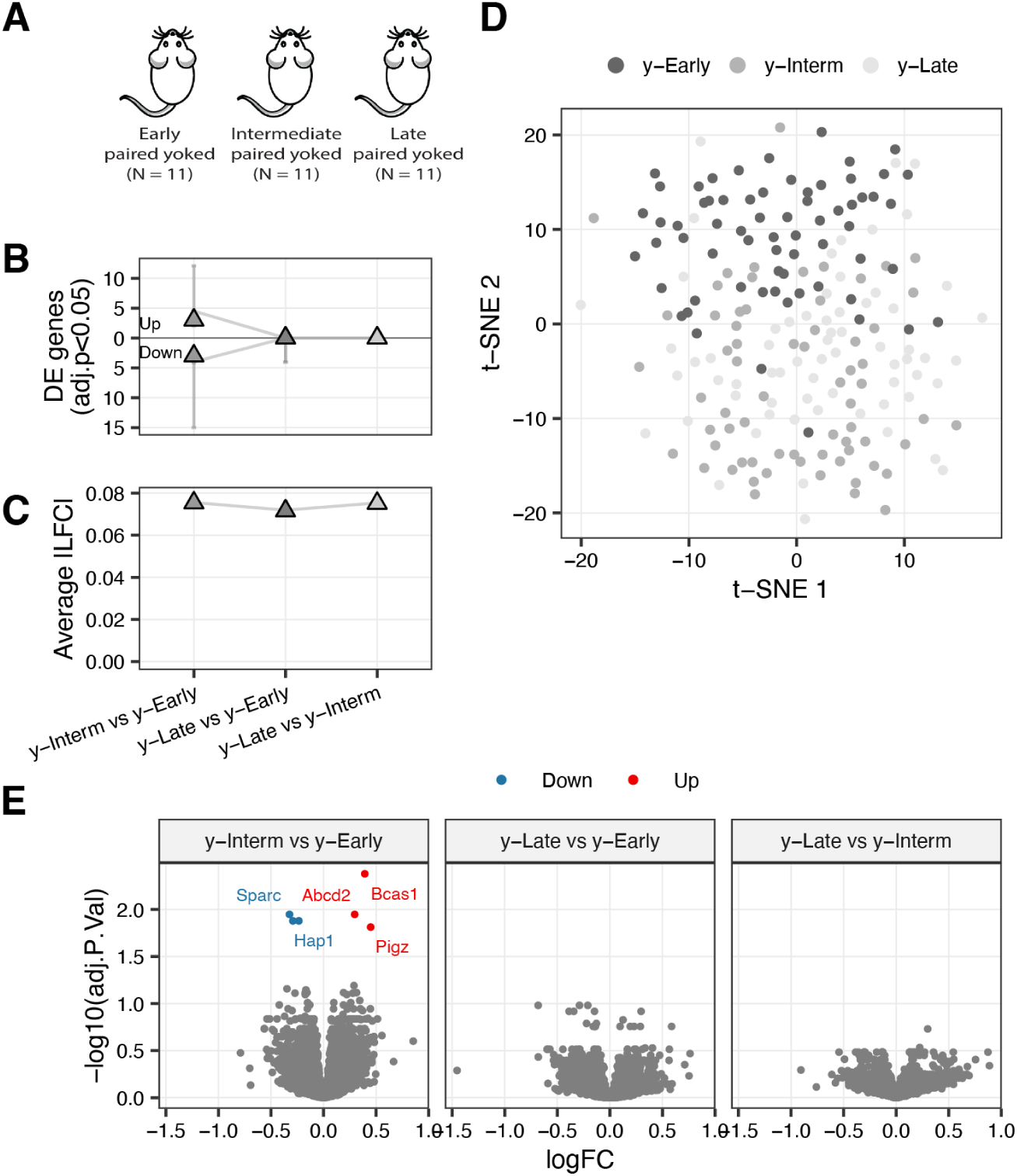
Changes in gene expression due to transition of the mice into the operant conditioning chambers. A - Schematic overview of the experimental setup regarding the yoked mice, who did not enter the operant conditioning chambers, while the yoked mice spent the same time in the operant conditioning chambers as the learners. B - The number of detected DE genes between each stage depicted in A. Mouse transition into the chamber induces few expression differences in the beginning. Total number of DE genes: y-Intermediate vs y-Early = 6; y-Late vs y-Intermediate = 0. C - The average effect sizes (|LFC|) across all genes between each stage depicted in A. B, C -Sampling distribution was generated using jackknifing, e.g., each mouse pair was removed once and the same model was refitted. Error bars: 95% percentile intervals. D - Dimensionality reduction using t-SNE based on the 100 most differentially expressed genes, ranked by their uncorrected p-values (PCs = 50, perplexity = 20, theta = 0.1). All random effect predictors and region effects (see methods) have been regressed out prior to clustering. E - Volcano plots depicting the observed log-fold changes and the adjusted p-values in each comparison. Blue and red dots indicate significantly down- and up-regulated genes, respectively (adjusted p-value<0.05).

**Extended Data Figure S8.**
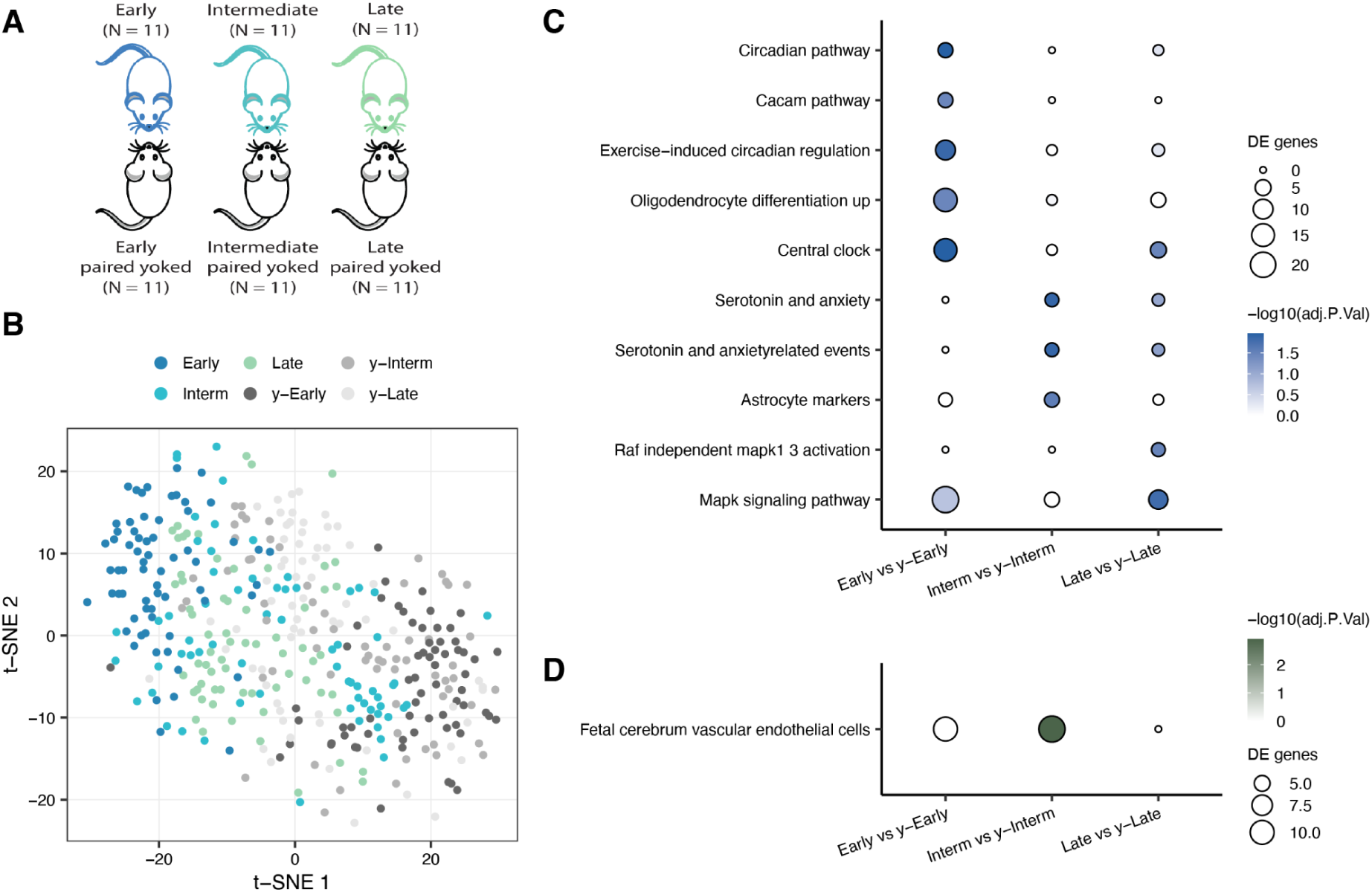
Additional analysis of the learning-specific transcriptional responses in the striatum. A – Schematic overview of the experimental setup regarding learning mice across time and the yoked paired mice. B – Dimensionality reduction using t-SNE based on the 100 most significant DE genes (PCs = 50, perplexity = 20, theta = 0.1). All random effect predictors and region effects (see methods) have been regressed out prior to clustering. Mice that had to learn cluster along the t-SNE 1 axis, distinguishing these from the yoked mice. C – Enriched M2 categories from selected published annotations on function in the mouse, when comparing learners with their respective yoked controls (BH-adjusted p-value<0.05)^44^. D – Enriched M8 categories from selected published annotations on cell type markers in the mouse, when comparing learners with their respective yoked controls (BH-adjusted p-value<0.05).

**Extended Data Figure S9.**
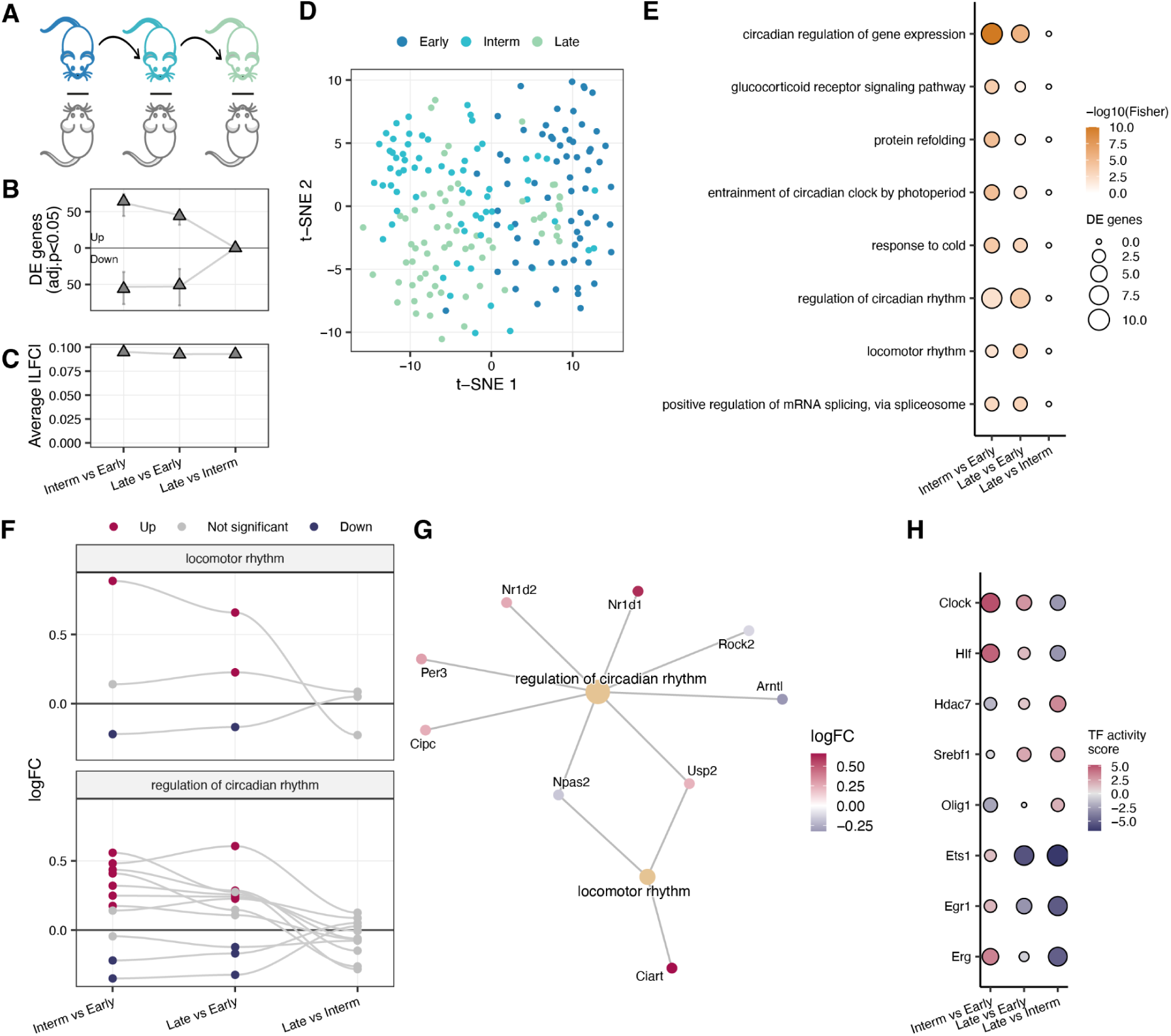
Expression differences between learning stages. A – Schematic overview of the experimental setup comparing the learning mice between different stages, while subtracting the across time and the yoked paired mice. B – The number of detected DE genes between adjacent learning stages, controlled for the non-learning effects using yoked mice. Total number of DE genes: Early vs Intermediate = 120; Late vs Early = 95, Intermediate vs Late = 0. C – The average effect sizes (|LFC|) across all genes of each adjacent learning stage, controlled for non-learning effects. B,C –Sampling distribution was generated using jackknifing, e.g., each mouse pair was removed once and the same model was refitted. Error bars: 95% percentile interval. D – Dimensionality reduction using t-SNE based on the 100 most significant DE genes (PCs = 50, perplexity = 20, theta = 0.1). All random effect predictors and region effects (see methods) have been regressed out prior to clustering. Mice from the Early learning stage cluster along the t-SNE 1 axis. E – Top five most significant gene ontology categories per comparison (Biological process, Fisher’s p-value<0.05). F – Expression differences of significant genes between learning stages from selected GO categories. G – Genes in pathways from F in the comparison between Late and Early learners. The nodes represent GO categories and the edges represent significantly DE genes belonging to the respective categories, colored according to their log-fold change. H – Predicted transcription factor (TF) activity differences between the adjacent learning stages, controlled for the non-learning effects, using the CollecTRI database. These associations were further pruned by the co-expression between the TF and the targets compared to TF and random genes (one-sided Wilcoxon rank sum test p-value<0.05).

**Extended Data Figure S10.**
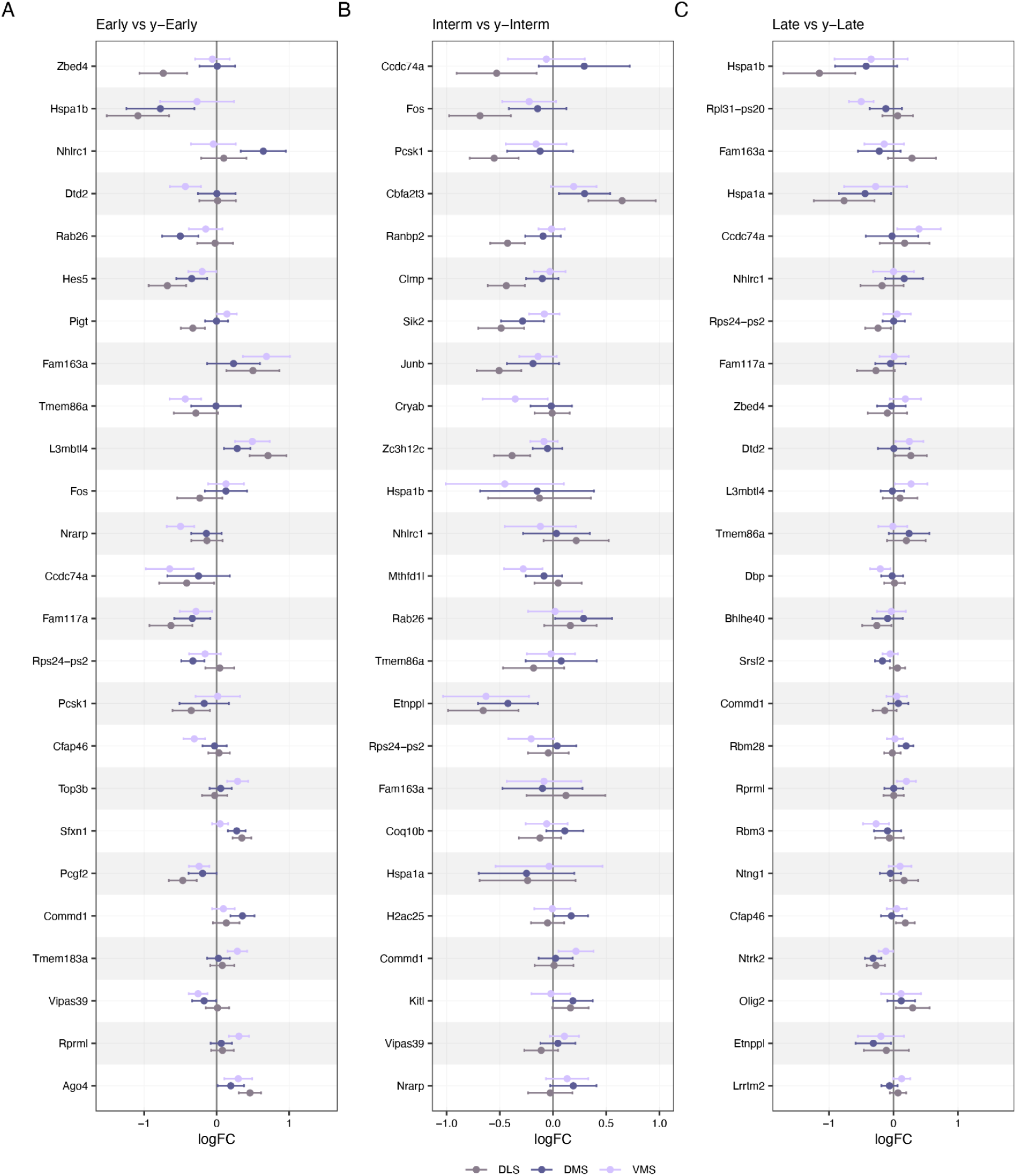
Top 25 genes with the largest log-fold change variation across striatal regions in response to learning. A – Early vs y-Early; B – Intermediate vs y-Intermediate; C – Late vs y-Late. VMS is depicted in light purple, DMS in dark blue, DLS in grey.

